# Estrogen remodels large-scale cortical dynamics to promote behavioral persistence

**DOI:** 10.64898/2026.10.25.761718

**Authors:** Meenakshi M. Asokan, Alissa Le, Dexter Tsin, Lucy Sirrs, Madeline Cramer, Stefan N. Oline, Forrest D. Rogers, Annegret L. Falkner

## Abstract

Sex-steroid hormones remodel the brain on slow timescales of hours to days^1–3^ to coordinate adaptive behaviors with changing bodily physiology^4–7^. Yet how this slow remodeling reorganizes and constrains neural computations that govern behavior on a moment-to-moment timescale remains poorly understood, particularly in cortical regions that support flexible social decision-making^8–11^. Here we combined longitudinal high-density neural recordings, quantitative behavioral phenotyping and estrogen-receptor manipulation in female mice across low and high estrogen states. Tracking behavior and neural dynamics using this dense sampling approach revealed that high estrogen increased signatures of behavioral and neural persistence. High estrogen reversibly increased male-directed social preference and behavioral persistence, effects that were abolished by estrogen receptor alpha knockdown in the medial prefrontal cortex. In the mPFC high estrogen produced more persistent neural states that were more closely aligned with ongoing behavior, increased state discriminability, and compressed within-state dimensionality. At the single-neuron level, estrogen increased tuning selectivity, lengthened intrinsic timescales, and stabilized social representations across days, even within the same neurons tracked longitudinally. High estrogen also strengthened recurrent connectivity during the resting state, exhibiting preferential connectivity between similarly tuned ensembles. Finally, incorporating these connectivity changes into a mean-field dynamical-systems model^12,13^ was sufficient to reproduce increased behavioral and neural-state persistence. Together, these findings show that estrogen oscillates cortical computations across multiday timescales, flipping the prefrontal cortex from flexible to persistent regimes that play a role in sustaining potentially costly social behaviors.

## INTRODUCTION

Across the lifespan, hormonal fluctuations repeatedly reshape the brain to meet changing physiological and behavioral demands^2,5–8,14–22^. Sex-steroid hormones such as estrogen orchestrate widespread structural and functional remodeling of the brain^4,23–26^, influencing motivation, affect, behavior and cognition. Although these hormones are broadcast throughout the body and brain, they can exert site-specific effects through the differential expression and action of hormone receptors^4,27–29^. One major mechanism is classical genomic signaling, in which estrogen binds to nuclear receptors that together act as transcription factors, which in turn regulate gene-expression programs influencing cellular excitability, synaptic connectivity, and circuit wiring^23,30^. In humans, large-scale structural changes accompany cyclic hormonal fluctuations in both sexes^3,31^, as well as more profound hormonal transitions such as pregnancy and perimenopause^18,19,32–34^. These periods of major hormonal change are also associated with increased vulnerability to social withdrawal and depression^35,36^, including during the postpartum and perimenopausal periods. Yet despite the pervasive effects of hormones on brain structure, function, and behavior, we still lack a unifying framework for how estrogen reorganizes neural dynamics, coding and computation to produce adaptive changes in behavior.

Mechanistic understanding of hormone-dependent neural remodeling has been built largely on foundational studies of subcortical hypothalamic circuits^5,7,14,16,22,37^, which are enriched in hormone receptors and govern components of social behaviors such as mating and parenting. These behaviors also require sensory integration, contextual evaluation, and moment-to-moment decision-making, processes that depend heavily on cortical circuits. Successful execution of these behaviors also depends on sustaining that choice and the underlying motivational state over time, demanding behavioral persistence rather than rapid switching between competing behavioral choices. The medial prefrontal cortex (mPFC) is a major hub for this top-down control of social behavior, with established roles in social recognition^10^, exploration^11^, and social decision-making^9^.

Although comparatively few studies have examined hormonal action in mPFC, existing work has shown hormone-dependent changes in dendritic spine density^17,38,39^ and in cell-type-specific mPFC projections to hypothalamic circuits that contribute to social preference^8^. Single-cell RNA sequencing has also revealed widespread expression of estrogen receptor alpha (ERα) across mPFC cell types and cortical layers^8^, further establishing the mPFC as a hormone-sensitive cortical region. However, there still exists a critical gap in understanding how hormone-dependent molecular and cellular changes are translated into population-level neural coding and dynamics in mPFC that support adaptive behavioral states.

Estrogen operates over slow timescales, with transcription-dependent effects unfolding over hours to days^1^, but most systems-neuroscience frameworks interrogate neural computation over seconds to minutes. This mismatch has made it difficult to determine how estrogen-dependent remodeling reorganizes the same neurons and networks across physiological states. In humans, dense-sampling approaches^3,31^ have begun to address this challenge by repeatedly imaging the same individual across endogenous hormonal transitions, revealing dynamic changes in functional network organization. Extending this logic to cellular-resolution measurements requires stable longitudinal approaches that can follow both population activity and individual neurons across days. Recent advances in chronic high-density electrophysiology^40,41^ and cross-session unit-tracking^42,43^ now make this tractable, providing a window into how hormonal state reshapes intrinsic neuronal properties, coding, and network organization over time.

Here, we hypothesized that estrogen shifts cortical computation toward a regime that prioritizes stability and persistence during high-estrogen states (when costly reproductive behaviors are most adaptive), whereas low-estrogen states favor greater flexibility and switching between behavioral choices. To test this, we combined longitudinal Neuropixels recordings with a simple direct-access social-choice assay in female mice with endogenous and experimentally induced estrogen cycles, together with deep behavioral phenotyping. We find that hormonal state does not simply modulate social preference, but shifts the dynamical regime of both behavior and cortical activity: high estrogen promotes persistent social choices and stable and rigid neural states, whereas low estrogen is associated with greater switching and flexibility. This shift is coordinated across computational scales, from more persistent and separable population representations to more selective and stable single-unit coding, longer intrinsic timescales, and increased clustered connectivity. Longitudinal tracking further reveals that these changes reflect dynamic reconfiguration of coding and connectivity within the same neurons across hormonal states rather than simply recruitment of different neural populations. Finally, estrogen preferentially strengthens recurrent connectivity in a structured manner within similarly tuned ensembles. These provide a network implementation for the shift toward stable population dynamics. Together, these findings establish a unifying computational framework that bridges network architecture, single-neuron coding, population dynamics, and behavior to explain how hormonal state shifts cortical circuits between flexible and persistent regimes.

## RESULTS

### Estrogen increases preference and persistence for male-directed interactions in females and requires mPFC receptor expression

One fundamental, normative function of estrogen-driven brain remodeling may be the prioritization of behaviors that are adaptive for a given physiological state. Consistent with this idea, circulating sex hormones tightly regulate social behaviors in rodents. Female mice are sexually receptive primarily during the peri-ovulatory estrus phase, when estrogen levels peak, and display predominantly rejective behaviors during other phases of the cycle^6,16,22^. Beyond regulating sexual receptivity, estrogen also modulates female preference for males^7^ and male-derived sensory cues^14^.

Here, we hypothesize that estrogen not only increases preference for males but also enables persistent neural and behavioral states that support sustained social choices that confer the most benefit during the reproductive window. To test this, we perform dense sampling of behavior and neural activity in both naturally cycling mice with endogenous hormonal cycles and ovariectomized mice with induced hormonal cycles **(Fig. 1a)**. In naturally cycling mice, although normal ovarian cycles involve coordinated fluctuations of both estrogen and progesterone, peak behavioral receptivity coincides with the peri-ovulatory estrogen surge and estrogen is considered to be the key regulator of social behavior. Moreover, mice can have variable cycles ranging from 4–7 days, and estrous staging based on vaginal cytology **(Extended Data Fig. 1a-b)** is a low resolution and noisy readout of the hormonal stage. Owing to the short cycle durations, it is therefore difficult to capture the precise low- and high-estrogen phases of the cycle, as well as obtain stable periods of low and high estrogen needed to study the underlying neural coding. To more precisely capture behavior and neural dynamics during high estrogen states, we induce controlled hormonal cycles by priming ovariectomized mice with daily injections of vehicle (oil) or 17β-estradiol (E2) to isolate and mimic the natural fluctuations of estrogen while also providing stable windows of low and high estrogen levels^6,14^. We densely sample behavior and neural activity via chronic Neuropixels implants, performing daily longitudinal recordings across low- and high-estrogen states during no-stimulus “resting-state” periods followed by the social-choice assay **(Fig. 1b)**. In the multi-day social-choice assay, the female subject can freely move and choose to interact with either a male or female conspecific tethered to the floor of one of two corners of the arena. This setup allows for direct social contact yet limits behavioral reciprocity from the conspecifics. We retain the same conspecifics and their locations in the arena across days to control for spatial coding, memory, and familiarity, and thereby isolate the effect of estrogen on social choice and the underlying neural computations.

**Fig. 1.**
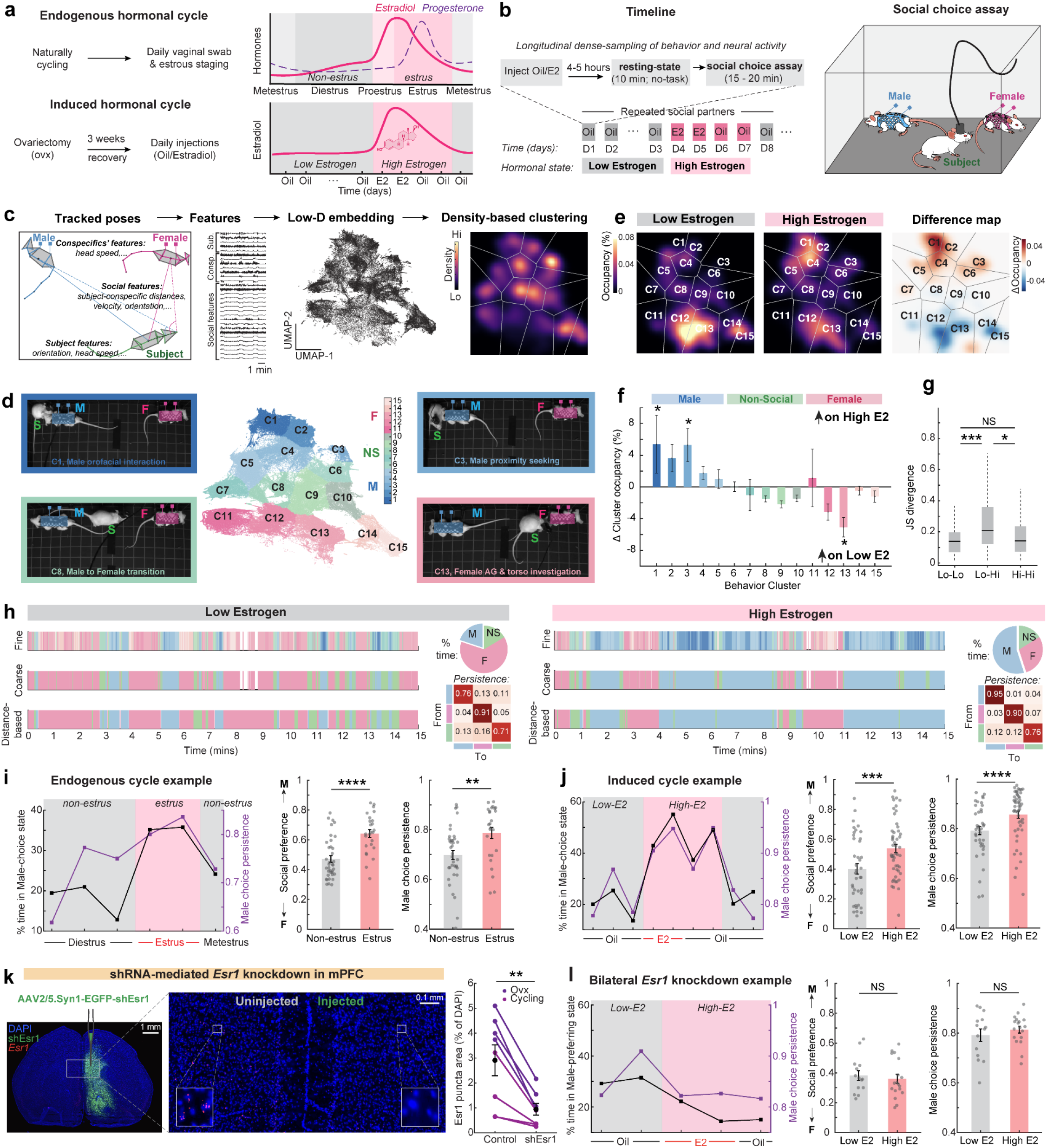
Estrogen receptor expression in the mPFC is necessary for estrogen-mediated increases in behavioral preference and persistence. **a,b,** Experimental timeline and schematic of endogenous and induced hormonal cycles (a), during which daily Neuropixels recordings were performed during resting state and the social-choice assay (b). E2, 17β-estradiol; F, female; Npx, Neuropixels. For naturally cycling mice, “non-estrus” includes metestrus and diestrus, whereas “estrus” includes proestrus and estrus. For OVX mice, “low E2” refers to baseline days with vehicle (oil) injection, and “high E2” includes the two days of E2 injection and the two immediately following days. **c,** Unsupervised behavioral quantification workflow, from tracked poses to frame-wise behavioral features, low-dimensional embedding using UMAP and density-based clustering to generate behavioral maps. **d,** UMAP embedding color-coded by coarse behavioral superclusters corresponding to male-directed (M; blue), female-directed (F; pink) and non-social (NS; green) interactions. Finer clusters (1–15) within each supercluster are shown using different shades of blue, pink and green. Representative video frames from a subset of clusters (1, 3, 8 and 13) are shown surrounding the embedding. **e,** Occupancy density maps in UMAP space averaged across low- and high-estrogen sessions for an example mouse, with the difference (high − low) shown on the right. Color scale, percentage of session time per bin (100×100 bin grid in UMAP space, Δ percentage points for the difference map); grey lines, cluster boundaries. **f,** Change in behavioral-cluster occupancy during high E2. Bars denote mean Δ occupancy during high E2 relative to each mouse’s mean low-E2 occupancy, expressed as percentage points of session time. (n = 6 mice, 24 high-E2 sessions; linear mixed-effects model accounting for repeated sessions within mice, with FDR correction across clusters.) **g,** Jensen–Shannon divergence between behavioral cluster-occupancy distributions for pairs of sessions grouped by hormonal state (low-low, low-high and high-high); 0 indicates identical occupancy distributions. Each point denotes one session pair. Box plots show the median and interquartile range. (n = 6 mice, with 32 low-low, 88 low-high and 36 high-high session pairs; within-mouse permutation tests with 10,000 shuffles of hormonal-state labels within each mouse, with FDR correction.) **h,** Behavioral ethograms from example low- and high-estrogen sessions showing fine behavioral clusters (top), coarse behavioral superclusters (middle) and comparable M/F/NS behavioral states defined using a supervised distance-based metric (bottom). Pie charts show the percentage of time spent in each behavioral state. Transition matrices show the probability of transitioning between states (off-diagonal) or remaining in the same state (diagonal) representing the choice persistence. **i,j,** Percentage of time in the male state (i) and male-choice persistence (j) across multiple days of the endogenous cycle (left) and induced cycle (right) in example mice. Summary plots show mean male preference and persistence during low and high E2 sessions; points denote individual sessions. (Endogenous cycle: n = 13 mice, 37/25 non-estrus/estrus sessions; induced cycle: n = 13 mice, 39/50 low/high E2 sessions; linear mixed-effects models accounting for repeated sessions within mice.) **k,** Viral-mediated local knockdown of estrogen receptor-α mRNA (*Esr1*) in mPFC using short-hairpin RNA (shEsr1). Unilateral injection was used to validate *Esr1* knockdown by RNAscope (left). *Esr1* puncta area, expressed as a percentage of DAPI area, was compared between the uninjected control and injected knockdown hemispheres (right). Each point denotes one mPFC section. (n = 2 mice, 4 sections per mouse; paired t-test.) **l,** Percentage of time in the male state and male-choice persistence in OVX mice with bilateral mPFC Esr1 knockdown, shown as in i,j. (n = 7 mice, 14/18 low/high E2 sessions; linear mixed-effects models accounting for repeated sessions within mice.) *Error bars indicate s.e.m. For all comparisons, *P < 0.05, **P < 0.01, ***P < 0.001, ****P < 0.0001. See Supplementary Table 1 for details of statistical tests and exact P values*.

To test whether female subjects display estrogen-state-dependent changes in behavior we first quantified moment-to-moment pose using unsupervised and distance-metric based analyses. We tracked the poses^44^ of all animals and converted the poses into a series of social and individual behavioral features **(Fig. 1c)**. To detect differences in the kinds of social interactions beyond simply assigning male versus female preference, we first took an unsupervised approach in which we generated a common low-dimensional behavioral embedding by performing non-linear dimensionality reduction of the high-dimensional behavioral features pooled across days and mice using an importance sampling approach^37,45,46^, followed by density-based clustering of the low-dimensional behavioral embedding. This approach led to a set of behavioral clusters that could be mapped to interpretable behaviors such as orofacial interactions, anogenital and torso investigations, proximity seeking, and transitions from one conspecific to another **(Fig. 1d)**. These fine-grained clusters also readily displayed a gradient from male-directed to non-social to female-directed behaviors. We grouped these fine-grained clusters into coarse-grained super-clusters that map onto male-directed, non-social, and female-directed behaviors using hierarchical clustering **(Extended Data Fig. 2a)**. To directly compare how the behavioral landscape changes between low- and high-estrogen states, we projected data from each low- and high-estrogen day onto the low-dimensional embedding, revealing broadly increased occupancy across male-directed clusters and reduced occupancy across female-directed clusters during high-estrogen days **(Fig. 1e-f)**. We found significant group differences in cluster occupancy across a subset of behavioral clusters, indicating that in the high-estrogen state, females spend more time performing male-directed orofacial interactions and proximity-seeking behaviors, and less time performing female-directed anogenital and torso investigation behaviors **(Fig. 1f)**. We observed a similar trend in endogenously cycling mice, where they displayed increased male proximity seeking and reduced female-directed orofacial and reciprocal interactions, as well as anogenital and torso investigation **(Extended Data Fig. 1c-d)**. In addition, there was greater behavioral similarity in terms of cluster occupancy between low-estrogen days and between high-estrogen days, whereas the two groups significantly differed from each other **(Fig. 1g)**.

Having established that high estrogen alters the behavioral landscape, we next asked whether it also changes the dynamics and persistence of male-directed interactions. To quantify male preference and persistence, we turned to a simple approach in which we quantified male-directed, female-directed, and non-social behaviors by thresholding based on two distance-based social features: the distance between the subject and each of the two conspecifics **(Extended Data Fig. 2c)**. We found a good match between the coarse behavior superclusters and the distance-based behavior states **(Fig. 1h, Extended Data Fig. 2c)**. We used the distance-based behavior state assignment to quantify the percentage of time spent in the male state and the transition probabilities between different states after binning the entire session at finer temporal resolution **(Fig. 1h)**. We then quantified social preference as the relative percentage of time spent in the male versus female state, and male-choice persistence as the transition probability of remaining in a male-directed state. We observed a reversible increase in social preference for males and male-choice persistence following the estrogen surge in mice with endogenous and induced cycles **(Fig. 1i-j, Extended Data Fig. 2d)**. Interestingly, while the overall percentage of time spent with female conspecifics decreased, female persistence remained unchanged, as did non-social state persistence **(Extended Data Fig. 2e-f)**.

Next, we tested whether local estrogenic action in the mPFC is critical for driving the hormone-dependent changes in social choice. Existing single-cell transcriptomic datasets indicate widespread but low-level expression of estrogen receptor alpha (ERα) across mPFC cell types and layers, with substantially lower expression of estrogen receptor beta (ERβ) and membrane-associated estrogen receptors such as GPER in the mPFC^8^. To test the role of ERα, we virally expressed eGFP-labeled short-hairpin RNA (shRNA) to knock down ERα expression bilaterally in mPFC **(Fig. 1k-l, Extended Data Fig. 3a-b)**. To validate ERα knockdown, we used fluorescent *in situ* hybridization to quantify *Esr1* in tissue sections taken from animals unilaterally injected with the shRNA in the mPFC **(Fig. 1k)**. Quantification of labeled probes showed a ∼66% reduction in *Esr1* in hemispheres expressing shRNA compared with uninjected control hemispheres in both naturally cycling and ovariectomized mice, despite an ∼3-fold upregulation of *Esr1* transcripts in the control hemispheres of ovariectomized mice **(Fig. 1k, Extended Data Fig. 3a)**. Estradiol priming in ovariectomized mice with bilateral ERα knockdown in mPFC failed to increase male preference or male-choice persistence, indicating that local estrogenic signaling via ERα in mPFC is critical for regulating hormone-dependent changes in social choice **(Fig. 1l, Extended Data Fig. 3c-e)**.

Overall, females showed a reversible increase in social preference for males following an estrogen surge in both estradiol-primed ovariectomized and naturally cycling mice, and elevated estrogen also increased behavioral persistence. These effects were abolished by local ERα knockdown in mPFC, indicating a critical role for local estrogenic signaling in regulating these behavioral changes.

### Estrogen increases neural-state persistence, improves population-level discriminability and compresses dimensionality in the mPFC

Having established that local estrogenic signaling in mPFC is critical for regulating hormone-dependent changes in social choice and behavioral persistence, we next sought to determine how changes in estrogenic state alter neural computations in mPFC to support these behavioral changes. We performed daily longitudinal, high-density recordings from hundreds of single units per session across mPFC subdivisions while subjects were placed in the arena during a no-stimulus “resting-state” window to record spontaneous activity and during the social-choice assay **(Fig. 1b)**. Recordings were performed using chronically implanted four-shank Neuropixels 2.0 probes^41^. Reconstruction of individual shank trajectories and recording sites confirmed that we precisely targeted mPFC subdivisions, with the majority of recorded units localized to prelimbic, infralimbic, or anterior cingulate cortex **(Fig. 2a-b, Extended Data Fig. 4a)**.

**Fig. 2.**
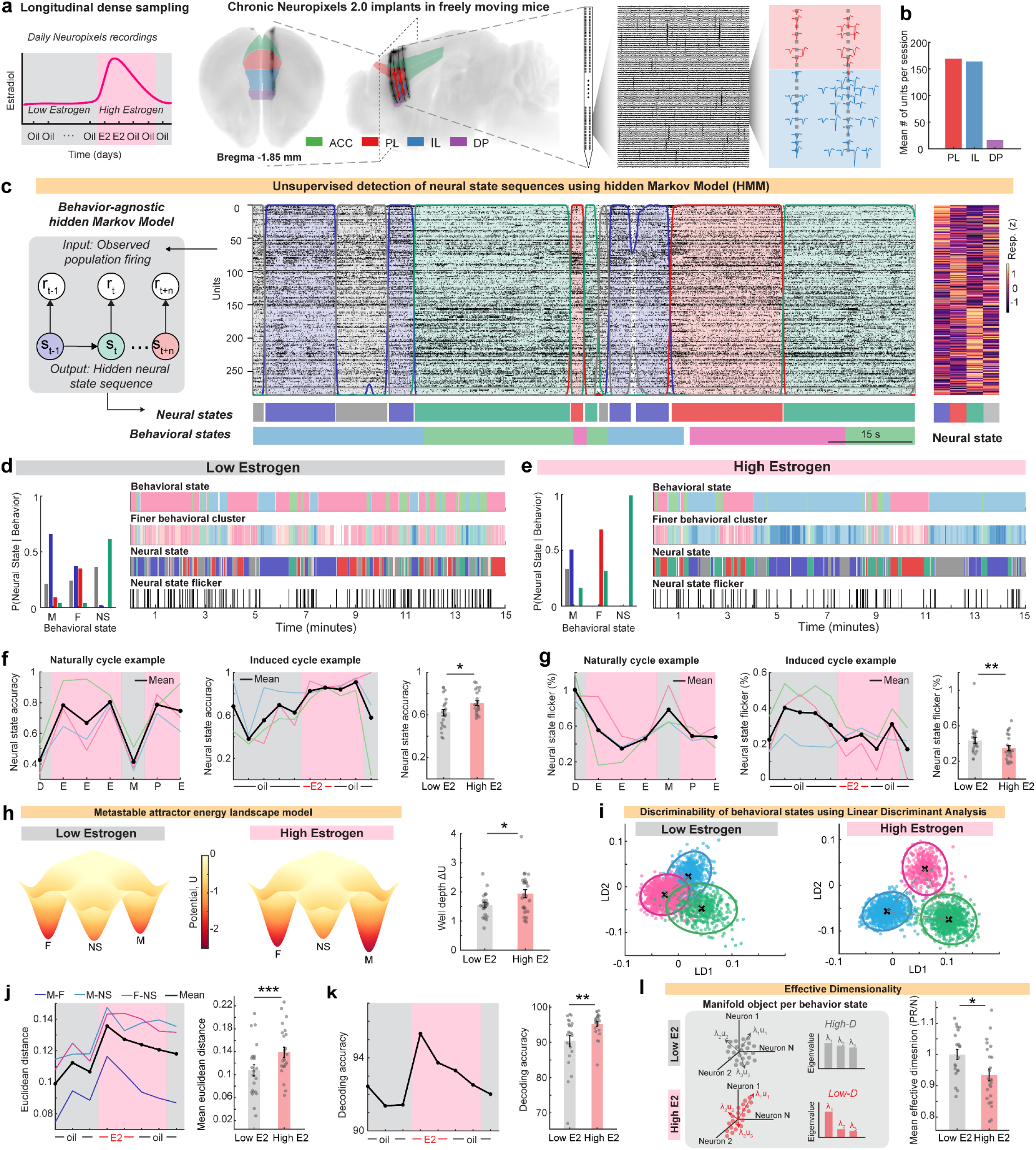
Estrogen increases neural-state persistence, improves population-level discriminability and compresses dimensionality in mPFC. **a,** Reconstruction of Neuropixels probe trajectories registered to the Allen CCF, showing anatomical targeting across mPFC subdivisions (left). Each line represents one shank of a four-shank Neuropixels 2.0 probe; red dots represent the 384 recording channels. Filtered voltage traces and example isolated single units spanning PL and IL (middle). **b,** Average single-unit yield per session across mPFC subdivisions (right). ACC, anterior cingulate cortex; PL, prelimbic cortex; IL, infralimbic cortex; DP, dorsal peduncular cortex. **c,** For each session, a hidden Markov model (HMM) (left, schematic) was fit to population spiking (r◻) throughout the social-choice assay, returning a sequence of hidden neural states (s◻), each corresponding to a pattern of population firing rates represented in the emission matrix (right). Center, population spiking raster during an example 2-min period, overlaid with latent-state posterior probabilities (colored curves); colored regions indicate intervals in which a state was detected with posterior probability >80%. Behavioral states, not used to fit the HMM, are shown below for comparison with the inferred neural states. **d-e,** Conditional probability P(neural state | behavioral state) used to establish a session-wise mapping between latent neural states and behavioral states (Left). Behavioral ethograms (Right) showing distance-based behavioral states and finer behavioral clusters, the corresponding neural states and neural-state flicker, are shown for example low E2 (d) and high E2 (e) sessions. **f,** Neural state accuracy defined based on alignment between behavioral states and matched latent state for each behavioral state and the mean across states (black) over consecutive days in an example naturally cycling (left) and induced cycling mouse (middle). Mean neural state accuracy across low E2 and high E2 sessions (right). (n = 6 mice, 22/24 low/high E2 sessions.) **g,** Neural-state flicker, expressed as a percentage of total frames, within each behavioral state and averaged across states (black) over consecutive days in an example naturally cycling (left) and induced cycling mouse (middle). Mean flicker across low E2 and high E2 sessions (right). (n = 6 mice, 22/24 low/high E2 sessions.) **h,** Potential-energy landscape U as an average across mice (left) and effective well depths (ΔU) estimated from median neural-state dwell times during corresponding behavior across low E2 and high E2 sessions (right). (n = 6 mice, 22/24 low/high E2 sessions.) **i,** Population activity projected into linear discriminant (LD) space during example low E2 (left) and high E2 (right) sessions. Each point represents population activity in one time bin, colored by the corresponding male-directed (blue), female-directed (pink) or non-social (green) behavioral state. **j,** Euclidean distances between behavioral-state representations for each pairwise comparison and the mean across comparisons (black) over consecutive days in an example mouse (left). Mean Euclidean distance across low E2 and high E2 sessions (right). (n = 6 mice, 23/24 low/high E2 sessions.) **k,** Support vector machine decoding accuracy of behavioral states from dimensionality-reduced population activity over consecutive days in an example mouse (left). Mean decoding accuracy across low E2 and high E2 sessions (right). (n = 6 mice, 23/24 low/high E2 sessions; linear mixed-effects model accounting for repeated sessions within mice.) **l,** Effective dimensionality of each behavioral-state manifold object (left, schematic). Mean effective dimensionality across manifolds, normalized within each mouse to its mean low E2 value (right). (n = 6 mice, 20/24 low/high E2 sessions.) *Points in summary plots denote individual sessions; bars and error bars indicate mean ± s.e.m. Low E2 and high E2 sessions were compared using linear mixed-effects models accounting for repeated sessions within mice. For all comparisons, *P < 0.05, **P < 0.01, ***P < 0.001, ****P < 0.0001. See Supplementary Table 1 for details of statistical tests and exact P values*.

To test the hypothesis that estrogen reorganizes population-level dynamics in mPFC to mediate changes in choice persistence, we segmented mPFC population activity during the social-choice assay in an unsupervised manner, agnostic to behavior, and identified corresponding neural-state sequences using a hidden Markov model (HMM)^47,48^. Using a separate model fit to each session, we inferred the sequence of hidden states through which population activity unfolds, with each state defined by a distinct activity pattern across the population, captured by the emission matrix **(Fig. 2c)**. Although we identified several fine-grained behavioral clusters, we saw that 3 - 4 latent states were sufficient for good model fit across days on the held-out data with minimal improvements from additional states **(Extended Data Fig. 5a-e)**. Importantly, the hidden states were inferred solely from population spiking activity and were completely naive to behavior, yet we observed reliable alignment between inferred neural states and behavioral states. Since HMMs were fit independently for each session, we mapped the session-specific latent neural states onto the three behavioral states (Male-directed, Female-directed, and Non-social) according to their behavioral occupancy, establishing a one-to-one correspondence between neural and behavioral states for each session. We then quantified neural state accuracy as the proportion of time within each behavioral state that the corresponding matched neural state was expressed. Neural state accuracy was significantly higher on high-estrogen days in both naturally cycling mice and ovariectomized mice with an induced estrogen surge **(Fig. 2d-f)**.

High estrogen not only increased the alignment between neural and behavioral states, but also reduced the tendency for neural activity to “flicker” between the hidden states while the behavioral state remains the same. We quantified neural-state flicker as a binary variable indicating whether the inferred neural state changed between consecutive time points within a fixed behavioral state. This measure reflects the moment-to-moment instability of neural population dynamics, with higher flicker indicating more frequent switching between inferred latent states. We observed a higher probability of neural-state flicker on low-estrogen days during ongoing behavior within all three behavioral states **(Fig. 2d-e,g, Extended Data Fig. 5f)**. In line with lower neural state accuracy and higher neural state flicker on low-estrogen days, we also found higher entropy in neural states within each behavioral state (**Extended Data Fig. 5g)**. There was no change in flicker rate aligned to transitions between fine-grained behavioral clusters **(Extended Data Fig. 5h)**, suggesting that the increased neural-state flicker observed under low estrogen was not simply driven by finer timescale behavioral transitions. Because HMMs can identify transitions in population activity that are not directly time-locked to external events or behavioral transitions, these results suggest that the stability of latent neural states itself varies with hormonal state. Prior work has interpreted such latent population states as metastable attractors that can be destabilized by internal noise or external perturbations^49–51^. Our findings suggest that estrogen increases the persistence of metastable neural states, stabilizing cortical population dynamics even when external stimuli and fine-grained behavior remain similar.

To develop a simple conceptual model for how estrogen reconfigures the dynamical landscape governing neural-state transitions, we next considered the inferred latent states within the metastable attractor framework. In this framework, metastable states are modeled as attractors of an effective dynamical system, with transitions between states occurring when fluctuations drive population activity out of one basin of attraction and into another^47,50^. We therefore used Kramers’ escape-rate theory^52^ to estimate effective attractor well depths, ΔU, from mean neural-state dwell times. High estrogen increased neural-state dwell times and correspondingly increased the estimated attractor well depths across behavioral states **(Fig. 2h)**. Within this framework, these results suggest that estrogen increases the stability of latent neural states and their resistance to noise-driven transitions.

Complementing the unsupervised identification of behaviorally aligned neural states, we next used a supervised classification approach to test whether estrogen alters the separability of population representations across behavioral states. To test this, we used linear discriminant analysis (LDA) and calculated the pairwise Euclidean distance between the latent-space neural representations of male-directed, female-directed, and non-social behavioral states. All pairwise distances increased following the estrogen surge, indicating more distinct and separable population representations of the three behavioral states **(Fig. 2i-j)**. Consistent with this increased separability, population-level decoding of behavior also improved under high estrogen **(Fig. 2k)**. Importantly, these changes were also reversible when estrogen levels returned to baseline **(Fig. 2j-k)**, and we observed similar reversible trends across multiple endogenous cycles in naturally cycling mice **(Extended Data Fig. 6a-b)**.

The increased separability of behavioral representations under high estrogen led us to ask whether this population-level reorganization was accompanied by a change in the geometry of neural population activity. Within a geometric framework, neural manifold objects represent population activity associated with different categorical states and occupy a low-dimensional subspace embedded within the higher-dimensional neural state space defined by the activity of individual neurons^53,54^. We therefore tested whether estrogen alters the geometric properties of these manifold objects, focusing on their effective dimensionality, which reflects the number of degrees of freedom needed to describe variance within each representation. For example, low-dimensional objects can be packed more efficiently, whereas high-dimensional objects can be less separable, constraining the representational capacity of the population^54,55^. To quantify this, we constructed manifold objects for each behavioral state and calculated the participation ratio as a measure of effective dimensionality. To standardize this measure across sessions and mice with different recorded population sizes, we analyzed randomly sampled neuronal subsets of a fixed size and repeated the analysis across resamples. High estrogen reduced the mean effective dimensionality of the behavioral-state manifold objects **(Fig. 2l)**. Thus, high estrogen was associated with both reduced dimensionality within behavioral representations and increased separation between them, consistent with a more compact population code.

Taken together, these results show that estrogen remodels mPFC population dynamics and geometry, stabilizing latent neural states with persistent dynamics while increasing the separability and compressing dimensionality of the behavioral representations.

### Estrogen increases selectivity and stability of social representations in mPFC single-units

We next tested whether changes to single-unit coding properties might underlie estrogen-mediated population-level reorganization. To assess how single units encode different behavioral states, we constructed rasters aligned to the onset of each event separately within the three behavioral states across the social-choice assay **(Fig. 3a-b)** and computed peri-event time histograms **(Fig. 3c)**. We quantified the evoked response as the area under the curve following state onset from the firing-rate fold change relative to a baseline window preceding state onset. Responses were then normalized, and each unit was assigned a response tuning (male-, female-, or non-social-tuned) based on its largest normalized response. To visualize tuning and response selectivity, each unit was projected onto a plane defined by a triangle intersecting the three Cartesian axes corresponding to its normalized responses to male-, female-, and non-social behaviors **(Fig. 3d)**.

**Fig. 3.**
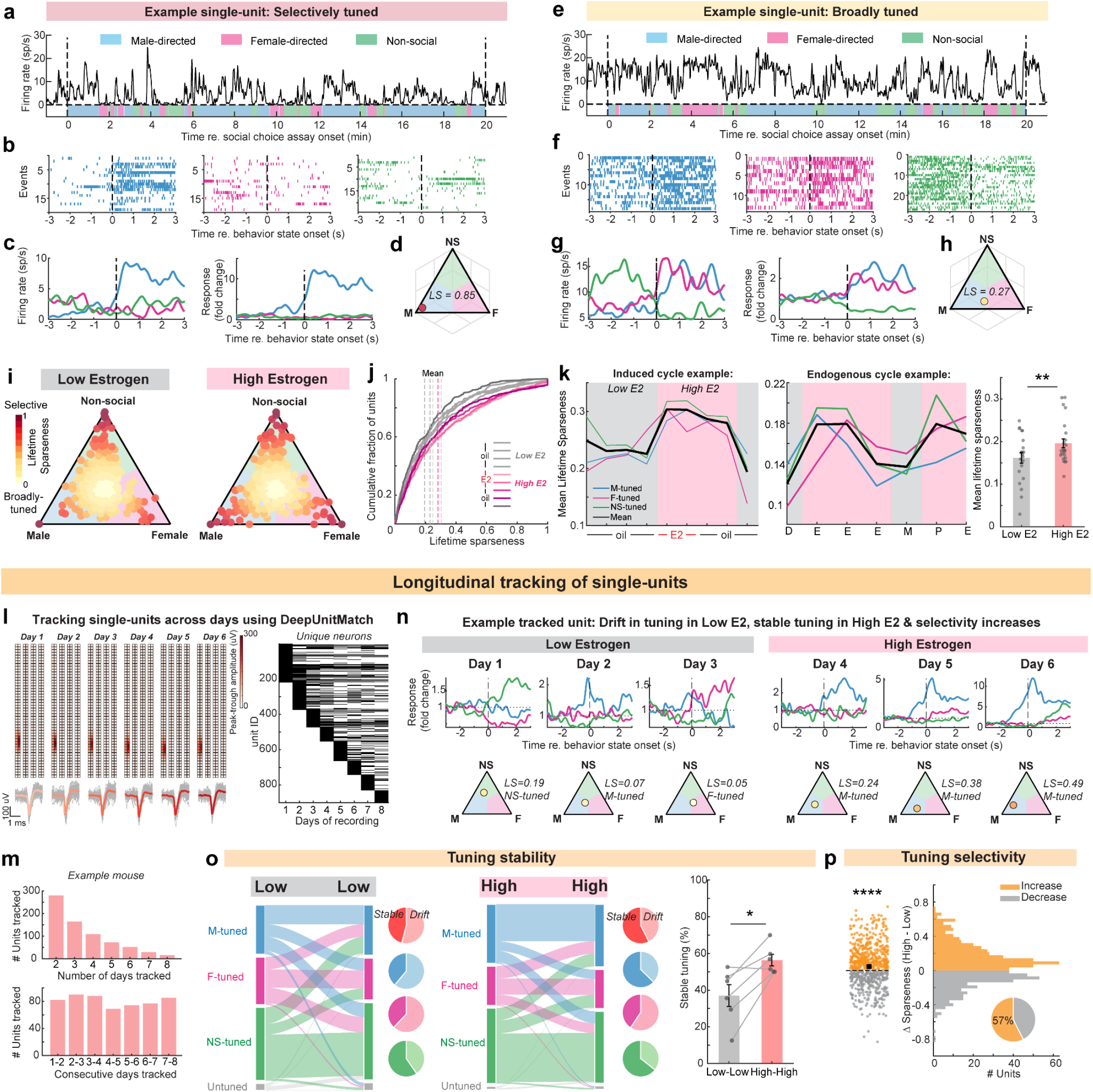
Estrogen increases single-unit tuning selectivity and stability. **a,** Firing-rate profile of an example single unit selectively tuned to male-directed interactions during the social-choice assay, with the corresponding behavioral ethogram shown below. **b,** Spike rasters for the single unit in a, aligned to the onset of each behavioral state. **c,** Peri-event firing-rate histograms averaged across events in b, shown as raw firing rate (left) and normalized fold change relative to a 1.5-s baseline window from −3 to −1.5 s before behavioral-state onset (right). **d,** Triangle projection depicting tuning and selectivity of the single unit in a-c, quantified using lifetime sparseness (LS). Each unit is projected onto an equilateral triangle with vertices at (0,0,1), (0,1,0), and (1,0,0), where the three axes correspond to the normalized responses to the three behavioral states. **e-h,** Same as a-d for an example single unit broadly tuned to both male and female behavioral states. **i,** Single units recorded during example low E2 (left) and high E2 (right) sessions projected onto the triangle as in (d); each point denotes one unit and is colored by lifetime sparseness. **j,** Cumulative distributions of lifetime sparseness across single units for each session from an example mouse; dashed lines indicate mean lifetime sparseness. **k,** Mean lifetime sparseness of units tuned to each behavioral state and the mean across tuning classes (black) over consecutive days in an example induced (left) and endogenously cycling mouse (middle). Mean lifetime sparseness across low E2 and high E2 sessions (right). (n = 6 mice, 21/24 low/high E2 sessions.) **l,** Amplitude profiles and waveforms of an example single unit tracked across six consecutive recording days (left). Tracking matrix showing the presence (black) of individual units across days, sorted by the first day detected (right). **m,** Number of units tracked across more than one day (left) and across consecutive pairs of days (right) in an example mouse recorded across eight consecutive days. **n,** Peri-event firing-rate histograms and triangle projections depicting tuning and selectivity (lifetime sparseness, LS) of example single units tracked across consecutive low E2 and high E2 days. The example unit shows tuning drift during low E2 followed by increased tuning stability and selectivity during high E2. **o,** Alluvial plots and pie charts from an example mouse showing the proportions of tracked units with stable versus drifting tuning across consecutive pairs of low E2 (left) and high E2 (middle) days. Percentage of units with stable tuning across low E2 and high E2 day pairs (right); points and connecting lines denote individual mice. (n = 6 mice; paired t-test on within-mouse differences.) **p,** Scatter plot and histogram show the change in lifetime sparseness between low E2 and high E2 after aggregating it for each tracked unit as the maximum value across days within each hormonal condition, and pie charts show the percentage of units with increased sparseness during high E2 (right). (n = 6 mice, 733 units; linear mixed-effects model accounting for units nested within mice.) *Points in summary plots denote individual sessions; bars and error bars indicate mean ± s.e.m. Low E2 and high E2 sessions were compared using linear mixed-effects models accounting for repeated sessions within mice. For all comparisons, *P < 0.05, **P < 0.01, ***P < 0.001, ****P < 0.0001. See Supplementary Table 1 for details of statistical tests and exact P values*.

mPFC is well known to contain neurons with “mixed selectivity,” in which single units exhibit broad tuning and respond to or encode multiple variables, particularly through nonlinear combinations^56,57^. Mixed selectivity has been associated with high-dimensional population representations and can support output flexibility and complex behaviors. On the other hand, “pure selectivity,” which refers to the selective response of a single unit to one variable, is associated with population-level dimensionality compression and allows for robust, high-fidelity execution of simple tasks^58,59^. Based on the need to prioritize reliable and persistent behaviors during high estrogen, together with our observation that estrogen reduces the effective dimensionality of behavioral-state manifold objects at the population level **(Fig. 2l)**, we hypothesized that estrogen would shift single-unit coding toward more selective responses. To test this, we computed Lifetime Sparseness (LS), a measure of how selectively a neuron responds across different variables. A unit with high LS responds selectively to a particular state and lies close to a vertex of the triangle **(Fig. 3a-d)**, whereas a unit with low LS responds more broadly across states and lies closer to the center of the triangle **(Fig. 3e-h)**. During high estrogen, a greater proportion of single units were positioned closer to the vertices of the selectivity triangle, accompanied by an increase in mean Lifetime Sparseness **(Fig. 3i-k)**. Interestingly, this increase in selectivity occurred in units tuned to all three behavioral states **(Fig. 3k, Extended Data Fig. 7a-c)**, while the proportion of units tuned to each state did not significantly vary across days or hormonal states **(Extended Data Fig. 7d-e)**. Thus, estrogen does not appear to recruit more neurons into a particular categorical tuning class, but instead sharpens the tuning of neurons within each class. The increase in selectivity was reversible when estrogen levels returned to baseline, and we observed a similar increase in tuning selectivity during estrus relative to non-estrus phases, independent of their order, in naturally cycling mice **(Fig. 3k, Extended Data Fig. 7c)**. Overall, these results suggest that single-unit coding during behavior shifts from broader tuning during low estrogen toward more selective tuning during high estrogen, consistent with a shift from a flexibility-prioritizing to a reliability-prioritizing coding regime.

To determine whether the increased selectivity and intrinsic timescales observed during high estrogen reflect recruitment of a different ensemble of neurons or reconfiguration of coding properties within the same neurons, we tracked single units longitudinally across days **(Fig. 3l-m)** using DeepUnitMatch^43^, a deep-neural-network based pipeline that matches neurons across sessions based on their spike waveforms following independent spike sorting of each recording session. This longitudinal approach crucially allowed us to distinguish population-composition changes from within-neuron reconfiguration across hormonal states. High-density probes such as Neuropixels enable stable chronic recordings from hundreds of neurons with high spatiotemporal resolution, while their geometry and density allow correction for electrode drift across days. Nevertheless, potentially because of residual electrode drift and other sources of recording variability, newly identified units appeared on each recording day **(Fig. 3l, right)**, and the number of units that could be continuously tracked decreased approximately exponentially with the number of recording days **(Fig. 3m)**. To specifically test whether broadly tuned neurons under low estrogen acquire more stable tuning during high estrogen, we stratified longitudinally tracked units by their baseline selectivity, defined as the mean Lifetime Sparseness across low-estrogen days. We focused this analysis on the low-selectivity group **(Fig. 3n)**, which showed broader responses across the three behavioral states under low estrogen and therefore provided greater dynamic range for detecting changes in tuning stability across hormonal states. Within this group, the proportion of units maintaining stable tuning was significantly higher between two high-estrogen days than between two low-estrogen days **(Fig. 3o)**. This remapping and drift in tuning during low estrogen may represent an additional feature of the broader, more flexible coding regime and has also been observed in hippocampal representations during low-estrogen phases in naturally cycling mice^2^. Other units maintained stable tuning across all days, but in both groups a greater proportion of single units increased their tuning selectivity, quantified by Lifetime Sparseness, during high estrogen **(Fig. 3p)**.

### Estrogen increases single-unit intrinsic timescales without altering spontaneous firing rates

We then turned to single-unit coding during “resting-state,” or periods of spontaneous activity in the absence of a stimulus or task. Intrinsic timescales of single units quantify the temporal persistence of neural activity at rest^60^ and have been proposed as a potential substrate shaping population dynamics and attractor states^61^. Intrinsic timescales can also vary dynamically with functional demands, for example increasing during working-memory maintenance^62^. To test whether intrinsic timescales increase during high estrogen in parallel with the more persistent population dynamics, we measured the intrinsic timescale of each single unit from the decay of its spike-count autocorrelation during spontaneous activity. Spontaneous activity was recorded during a 10-minute period at the beginning of each session, with spikes binned in 5-ms time bins across the entire window. Intrinsic timescales are typically estimated by fitting the autocorrelation with an exponential decay function, which appears as a straight line in semilogarithmic coordinates, with a slope determined by the inverse of the timescale. To capture multiple potential timescales in single-neuron dynamics, we fitted each autocorrelation with linear mixtures of exponential decay functions and estimated an effective decay time constant **(Fig. 4a-b)**. For each neuron, we fitted models containing one to four timescale components (M = 1-4) and selected the optimal number using the Bayesian information criterion. Only units with a good model fit, comprising ∼50% of units across sessions, were included in subsequent timescale analyses **(Extended Data Fig. 8a)**. Consistent with prior studies^60^, many neurons were best fit by models containing two or three exponential components, and interestingly, a significantly higher proportion of units displayed best fits with higher number of timescales during high-estrogen, suggesting the presence of multiple timescales within the activity of individual neurons **(Extended Data Fig. 8b)**. Consistent with our hypothesis, a greater proportion of neurons displayed longer intrinsic timescales, and the median intrinsic timescale across the population increased during high estrogen **(Fig. 4c-d)**. The increase in intrinsic timescales was not accompanied by a change in spontaneous firing rates, and the spontaneous firing-rate distributions and the median spontaneous firing rate did not vary with estrogenic state **(Fig. 4e-f)**. We also observed a similar increase in intrinsic timescales during estrus relative to non-estrus phases in naturally cycling mice **(Extended Data Fig. 8c)**. Estrogen also increased intrinsic timescales within longitudinally tracked single units **(Fig. 4g-h, Extended Data Fig. 8d)** while spontaneous firing rates remained stable across days **(Fig. 4i)**.

**Fig. 4.**
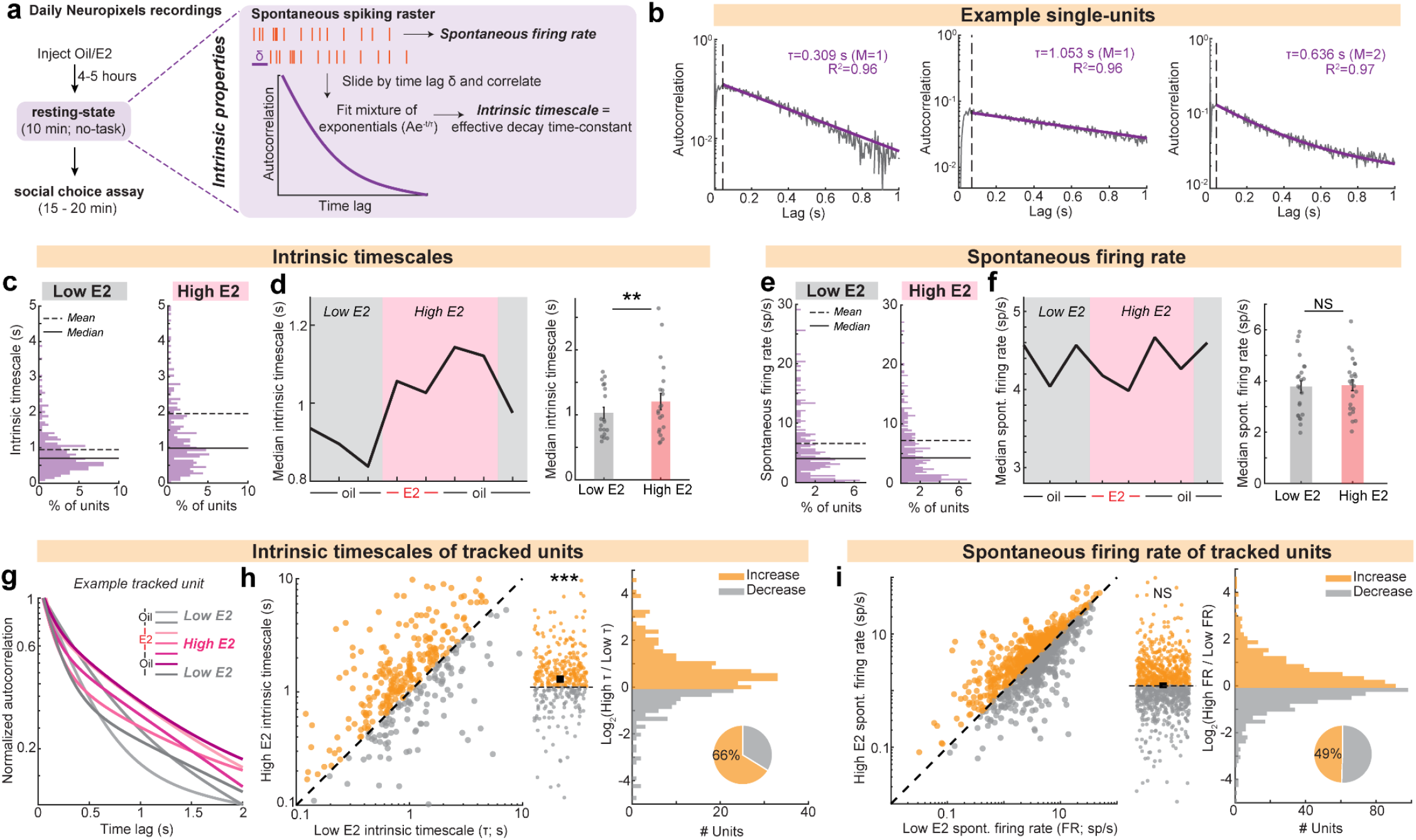
Estrogen increases single-unit intrinsic timescales without altering spontaneous firing rates. a, Schematic of intrinsic-timescale estimation for individual single units, defined as an effective decay time constant obtained by fitting a mixture of exponentials to the autocorrelation of spontaneous spiking activity. **b,** Autocorrelation data (grey) and corresponding mixture-of-exponentials fits (purple) for example single units. M denotes the number of exponential components in the best-fitting model. **c,** Distribution of intrinsic timescales during example low E2 and high E2 sessions from an example mouse. **d,** Median intrinsic timescale over consecutive days in an example mouse (left). Median intrinsic timescale across low E2 and high E2 sessions (right). (n = 6 mice, 19/21 low/high E2 sessions.) **e,** Distribution of spontaneous firing rates during example low E2 and high E2 sessions from an example mouse. **f,** Median spontaneous firing rate over consecutive days in an example mouse (left). Median spontaneous firing rate across low E2 and high E2 sessions (right). (n = 6 mice, 23/24 low/high E2 sessions.) **g,** Autocorrelation data and mixture-of-exponentials fits for an example single unit tracked across consecutive low E2 and high E2 days. **h,** Intrinsic timescales of single units tracked across at least one low E2 and one high E2 day (left). For each unit, the intrinsic timescale was averaged across days within each hormonal condition. Scatter plot and histogram show the change in intrinsic timescale between low E2 and high E2, and pie charts show the percentage of units with increased timescales during high E2 (right). (n = 6 mice, 355 units; linear mixed-effects model fit to log₂-transformed values, accounting for units nested within mice.) **i,** Spontaneous firing rates of single units tracked across at least one low E2 and one high E2 day (left). For each unit, spontaneous firing rate was averaged across days within each hormonal condition. Scatter plot and histogram show the change in spontaneous firing rate between low E2 and high E2, and pie charts show the percentage of units with increased firing rates during high E2 (right). (n = 6 mice, 984 units; linear mixed-effects model fit to log₂-transformed values, accounting for units nested within mice.) *For all comparisons, *P < 0.05, **P < 0.01, ***P < 0.001, ****P < 0.0001. See Supplementary Table 1 for details of statistical tests and exact P values*.

Taken together, we see that estrogen reconfigures single-unit coding from a regime characterized by broader tuning, greater representational drift, and shorter intrinsic timescales toward one characterized by more selective and stable tuning and longer intrinsic timescales. These changes are consistent with a shift from a coding regime that prioritizes flexibility during low estrogen toward one that promotes persistence and stability during high estrogen, even for epochs outside social behavior.

### Estrogen strengthens recurrent connectivity within similarly tuned mPFC ensembles

We next asked what circuit-level changes could implement the observed reorganization of neural coding and population dynamics. Given that local ERα signaling in mPFC is required for estrogen-dependent changes in social choice, one plausible mechanism is a change in local connectivity. ERα can act through classical genomic signaling to regulate gene-expression programs related to wiring and synaptic coupling^23,30^. In support of this, estrogen-dependent changes in dendritic spine density have been reported across several brain regions, including hippocampus^2^ and the mPFC^17,39^. We therefore tested whether estrogen alters connectivity within the local mPFC network.

To estimate connectivity within the local network, we computed resting-state spike cross-covariance for each pair of single units during the 10-minute spontaneous-activity period recorded at the beginning of each session. Because it is inherently difficult to distinguish synaptically connected pairs from pairs receiving common input from a third neuron, particularly in densely and recurrently connected regions such as mPFC, we applied stringent inclusion criteria to identify pairs as connected^63^. We classified a pair as connected if it displayed significant cross-covariance exceeding the confidence interval within a ±5-ms time lag, excluding zero lag **(Fig. 5a-b)**. Because cross-covariance was computed at rest, in the absence of a task, social partner, or stimulus, the influence of time-locked external drive that could force multiple neurons to fire together should be minimized, increasing the contribution of local network interactions to the observed correlated firing. Excluding pairs with a cross-covariance peak at zero lag further reduced the contribution of pairs dominated by shared common input. Moreover, the short-lag 5-ms threshold enriched for direct or near-direct synaptic interactions. Cortical excitatory monosynaptic connections have been reported to exhibit delays of approximately 1 to 4 ms^63^, whereas longer-latency interactions are more likely to reflect indirect pathways. Consistent with this choice, the empirical distribution of significant cross-covariance peak lags showed a sharp drop around 5 ms, beyond which the distribution approached a relatively flat baseline **(Extended Data Fig. 9a)**. Pairs that passed these criteria displayed a characteristic short-latency asymmetric cross-covariance profile with a rapid rise followed by a slower decay^63,64^ **(Fig. 5a)**.

**Fig. 5.**
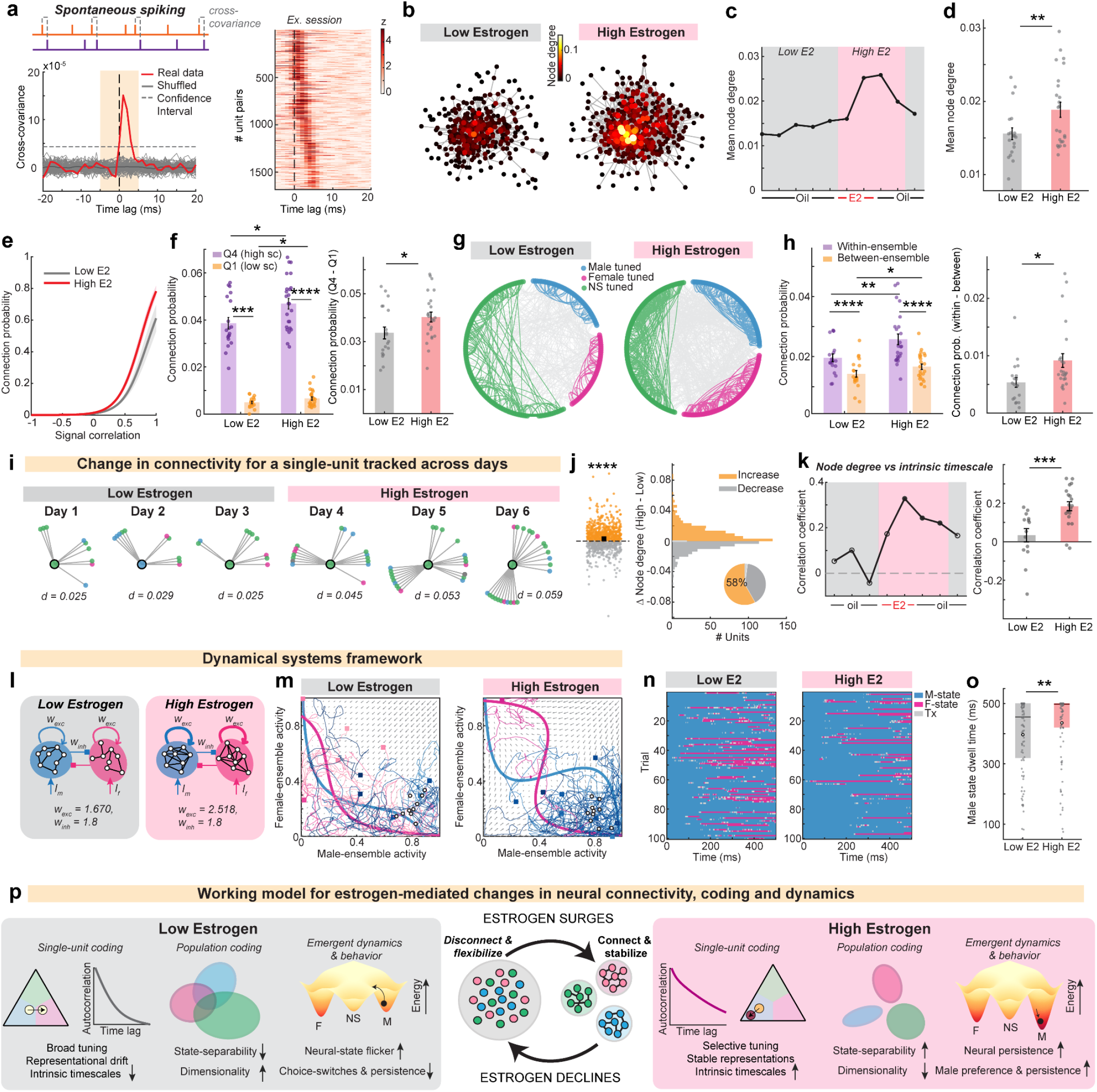
Estrogen strengthens recurrent connectivity in mPFC between similarly tuned neurons. **a,** Schematic and example spike-time cross-covariance for a pair of single units during spontaneous activity (left), and normalized cross-covariance for all significantly connected unit pairs from an example session (right). Significant connections were defined by cross-covariance exceeding the confidence interval at a non-zero |lag| ≤ 5 ms. **b,** Network graphs depicting resting-state functional connectivity within the mPFC network during example low E2 (left) and high E2 (right) sessions. Nodes denote single units and edges denote significant cross-covariance as defined in a. Node color indicates normalized node degree, defined as the number of connections to each node divided by the total possible number of connections for that node. **c-d,** Mean normalized node degree over low E2 and high E2 recording days in an example mouse (c) and summary across all low E2 and high E2 sessions (d). (n = 6 mice, 19/24 low/high E2 sessions.) **e,** Connection probability as a function of signal correlation (sc) between unit pairs, fit using logistic regression for an example mouse. Curves show mean fits across sessions and shaded regions indicate s.e.m. **f,** Mean connection probability for unit pairs in the lowest (Q1; low sc) and highest (Q4; high sc) quartiles of signal correlation across low E2 and high E2 sessions (left) and difference between Q4 and Q1 connection probability for all sessions (right). (n = 6 mice, 19/24 low/high E2 sessions.) **g,** Chord plots from example low E2 (left) and high E2 (right) sessions. Dots denote single units grouped by tuning, and arcs denote significant connections as defined in (a). Within-ensemble connections are shown in the corresponding tuning-group color and between-ensemble connections in grey. **h,** Connection probability for within-ensemble and between-ensemble unit pairs across low E2 and high E2 sessions (left) and difference between within-ensemble and between-ensemble connection probability for all sessions (right). (n = 6 mice, 19/24 low/high E2 sessions.) **i,** Day-by-day connectivity of an example single unit tracked across consecutive low E2 and high E2 days. The tracked unit is shown at the center, with significantly connected units on the surrounding ring. Connected partners are ordered by the number of days on which they appear and positioned anticlockwise from the right. **j,** Normalized node degree of single units tracked across at least one low E2 and one high E2 day (left), averaged across days within each hormonal condition. Scatter plot and histogram show the change in normalized node degree between low E2 and high E2, and pie charts show the percentage of units with increased node degree during high E2 (right). (n = 6 mice, 943 tracked units; linear mixed-effects model accounting for units nested within mice.) **k,** Spearman’s correlation coefficient between normalized node degree and intrinsic timescale over low E2 and high E2 recording days in an example mouse (left; filled circle, p < 0.05; open circle, p ≥ 0.05) and summary across all low E2 and high E2 sessions (right). (n = 5 mice, 14/19 low/high E2 sessions.) **l,** Biophysical model of social-choice dynamics incorporating the proposed circuit changes from low E2 (left) to high E2 (right). Each node represents a population tuned to male (blue) or female (pink) behavioral states, with recurrent self-excitation and mutual inhibition between populations. **m,** Phase-space representation of male- and female-tuned ensemble activity in the model. Arrows in the velocity vector field show the direction of flow, with arrow size indicating the rate of change; blue and pink thick lines indicate the nullclines for the male- and female-tuned populations, respectively. Example trajectories are initialized with greater activity in the male-tuned ensemble (open circles), with endpoints of the 500 ms trial indicated by filled squares. Trajectories ending in the female basin of attraction are shown in pink. **n,** Simulated neural-state sequences from 100 trials as in (m). Neural states are assigned as male or female when trajectories occupy the corresponding basin of attraction and as grey when trajectories lie at the boundary between basins. **o,** Male neural-state dwell times across simulated low E2 and high E2 trials. (n = 100 simulated trials; two-sided Wilcoxon rank-sum test; box plots show the median, inter-quartile range and open circle denotes the mean) **p,** Working model of estrogen-mediated changes across levels of organization, linking functional connectivity, single-unit and population coding, emergent neural dynamics and behavior. *Low E2 and high E2 sessions in d, f, h and i were compared using linear mixed-effects models accounting for repeated sessions within mice, with FDR correction for f, h. For all comparisons, *P < 0.05, **P < 0.01, ***P < 0.001, ****P < 0.0001. See Supplementary Table 1 for details of statistical tests and exact P values*.

To visualize differences in network connectivity between high and low estrogen states, we used graph-theoretic representations in which each single unit was depicted as a node, each significant connection as an edge, and node color represented normalized node degree, defined as the number of direct connections to a node normalized by the total number of possible connections for the recorded population in that session. High estrogen increased both the density of connections and the mean normalized node degree of the population **(Fig. 5b-d)**. Importantly, node degree was relatively stable across multiple low-estrogen days, increased during high estrogen for units tuned to all three behavioral states, and returned toward baseline 2-3 days after the second estradiol dose as estrogen levels declined **(Fig. 5c, Extended Data Fig. 9b-c)**. We observed a similar oscillatory modulation of node degree across multiple endogenous hormonal cycles in naturally cycling mice **(Extended Data Fig. 9d)**. In contrast, for unit pairs exhibiting significant negative cross-covariance, we did not observe a significant estrogen-dependent change in node degree **(Extended Data Fig. 9e-h)**.

Cortical networks are often organized into functional subnetworks with greater within-cluster than between-cluster connectivity, and such like-to-like connectivity motifs have been described across several cortical areas^63,65^. Clustered recurrent architectures can support metastable dynamics^52^, and theoretical work has also linked increased recurrence to lower-dimensional population representations and more robust downstream readout^66^. These observations led us to hypothesize that estrogen preferentially increases recurrent connectivity within similarly tuned ensembles rather than uniformly increasing connectivity throughout the network. To test this, we first computed signal correlation between neuronal pairs measured during behavior and compared this to resting-state connection probability **(Fig. 5e)**. We observed that pairs with high signal correlation, indicative of similar behavioral tuning, displayed a higher probability of connectivity during the resting state (outside of behavior), revealing a like-to-like connectivity motif within the mPFC network **(Fig. 5e-f)**. High estrogen increased connection probability among pairs with high signal correlation **(Fig. 5e-f)**, suggesting that the overall increase in network connectivity is concentrated within similarly tuned ensembles. To test this more directly, we treated neurons tuned to the three behavioral states as three subnetworks and compared within- and between-cluster connectivity. High estrogen increased within-ensemble recurrence across all three subnetworks **(Fig. 5g-h)**. Within-ensemble connectivity exceeded between-ensemble connectivity in both low- and high-estrogen states, and the difference in within-ensemble versus between-ensemble connection probability was higher in high-estrogen states **(Fig. 5h)**. For unit pairs exhibiting negative cross-covariance, we observed the opposite organizational motif: connection probability decreased as a function of signal correlation **(Extended Data Fig. 9k)**, and between-cluster connectivity exceeded within-cluster connectivity in both low- and high-estrogen states without significant estrogen-dependent changes **(Extended Data Fig. 9i-m)**. To determine whether the estrogen-dependent increase in connectivity reflected recruitment of a different recurrent ensemble or remodeling of connectivity within the same neurons, we compared normalized node degree in single units tracked longitudinally across days. The same tracked neurons displayed higher normalized node degree during high-estrogen phases **(Fig. 5i-j)**, indicating that estrogen-dependent network densification can arise through within-neuron reconfiguration rather than solely through changes in population composition.

Increased recurrent connectivity is predicted to stabilize local activity^52^ and prolong temporal integration, so we next asked whether the neurons that are more connected during high estrogen also exhibited longer intrinsic timescales. We computed correlations between normalized node degree and intrinsic timescale across single units within each session and found a significant positive relationship on high-estrogen days but not on low-estrogen days **(Fig. 5k, Extended Data Fig. 10a-c)**. This state-dependent relationship suggests that the increase in resting-state connectivity during high estrogen is preferentially associated with neurons exhibiting longer intrinsic timescales. In contrast, we did not observe significant correlations between Lifetime Sparseness and node degree or between Lifetime Sparseness and intrinsic timescale **(Extended Data Fig. 10a-c)**.

Finally, we used these empirically observed estrogen-mediated changes in structured mPFC connectivity to simulate whether the observed estrogen-dependent changes in network architecture could generate more persistent neural dynamics with reduced neural-state flicker. To do this we implemented a compact mean-field mutual-inhibition dynamical-systems model commonly used to study decision-making dynamics in mPFC^12,13,67^. In the model, male- and female-tuned excitatory ensembles were each endowed with recurrent self-excitation (w_exc_), and coupled through effective mutual inhibition (w_inh_), **(Fig. 5l)**. Each excitatory pool was modeled as engaging a shared inhibitory population that provided broad feedback inhibition, thereby generating competition between the two excitatory ensembles. Owing to the mean-field approximation, the state of the system at each moment was described by the average activity of neurons within each ensemble. To model a local epoch of male-directed behavior, we applied a small constant input to the male-tuned ensemble. Using the empirically observed difference in within-ensemble connectivity as a reference, we parameterized w_exc_ and w_inh_ for low- and high-estrogen conditions such that both remained within a bistable regime **(Fig. 5l)**. We then simulated repeated trials initialized with greater activity in the male-tuned than female-tuned ensemble and introduced small random perturbations to test the stability of the resulting trajectories **(Fig. 5m)**. Under the low-estrogen parameterization, trajectories were more susceptible to crossing the boundary between the two basins of attraction and switched between the two states **(Fig. 5m,n)**. In contrast, under the high-estrogen parameterization, trajectories remained more persistently within the initial basin of attraction, producing fewer state transitions and longer dwell times **(Fig. 5o)**. Thus, incorporating the estrogen-dependent changes in recurrent self-excitation into the model while keeping the mutual inhibition constant was sufficient to recapitulate key features of the experimentally observed HMM-based neural-state dynamics.

Taken together, these results indicate that an estrogen surge remodels mPFC network architecture by increasing overall connectivity while preferentially strengthening connectivity within similarly tuned ensembles. Incorporating these connectivity changes into a dynamical-systems model reproduced the experimentally observed increase in neural-state persistence and reduction in state flicker, providing a network-level framework linking estrogen-dependent circuit reconfiguration to changes in population dynamics.

## DISCUSSION

A central advance of this study is to link hormone-dependent changes in network organization, single-neuron coding, population dynamics, and behavior within a common computational framework. Our results suggest that the estrogen-dependent increase in clustered connectivity provides a plausible network substrate for longer intrinsic timescales and more selective and stable single-neuron representations. At the population level, these changes are accompanied by reduced neural-state flicker, longer state dwell times, greater behavioral-state separability, and lower effective dimensionality. Within an attractor framework, these observations are consistent with a shift toward more stable population states that are less susceptible to fluctuation-driven transitions. Conversely, broader tuning, greater representational drift, overlapping population representations, and shorter intrinsic timescales during low-estrogen states may support a more flexible regime in which neural and behavioral states switch more readily. Thus hormonal state appears to coordinate multiple features of cortical computations along a broader flexibility-persistence axis **(Fig. 5p)**.

These findings across biological scales are functionally interrelated. At the molecular level, mPFC expresses nuclear estrogen receptor alpha which regulates genes involved in connectivity^8,30^. At the network level, we not only see an increase in the density of connections within mPFC but also clustered connectivity within similarly tuned ensembles which have been proposed to stabilize metastable dynamics by deepening attractor states^52^, while also reducing dimensionality^66^ and supporting robust downstream readout. At the single-neuron level, selective tuning may emerge from clustered connectivity and has been associated with reduced population dimensionality^59^ and robust, high-fidelity performance on simpler tasks^58^. We also find a positive relationship between node degree and intrinsic timescales under high-estrogen conditions, suggesting that neurons with greater connectivity may exhibit slower temporal dynamics. Intrinsic timescales have, in turn, been proposed as a potential substrate for persistent population activity^61^. Finally, at the population level, deeper attractor wells and reduced dimensionality may support more separable representations, allowing distinct representations to be encoded more efficiently within the same population space^54,57^ and potentially supporting persistent behavioral states. Together, these findings provide a unifying understanding of how hormonal state reorganizes cortical computations to support adaptive behavior.

These system-level changes observed in mPFC likely arise from local estrogenic signaling at slow genomic timescales. The level of expression of ERα in mPFC is higher than membrane-associated estrogen receptors such as GPER^8^. These nuclear receptors directly regulate neural gene expression through interactions with genomic regulatory elements such as the estrogen response elements, accompanied by changes in chromatin accessibility^30^. In the adult female brain, ovarian hormone fluctuations are associated with dynamic changes in chromatin organization and transcriptional activity^68,69^. Recent single-cell multiome profiling further suggests that chromatin accessibility and gene expression are dynamically regulated across reproductive transitions^20^, raising the possibility that chromatin remodeling also primes the genome for subsequent transcriptional changes during future or sustained hormone exposure. Together with our finding that ERα in the mPFC is critical for estrogen-dependent modulation of social choice, these studies raise the possibility that hormone-dependent transcriptional and chromatin changes contribute to the systems-level reorganization observed here. We propose that this molecular action leads to changes in local recurrent connectivity that could in turn be a plausible underlying mechanism for coordinating changes in neural persistence and selective, stable tuning. In addition to that, estrogen-dependent changes in cell-type-specific excitability^5,8^, neuromodulatory tone^15,70^, and activity in upstream brain regions^21,71^ may also contribute. We also cannot rule out additional contributions from faster, non-genomic estrogen signaling^72^, and the interplay between genomic and non-genomic mechanisms may contribute to differences in the timescales over which distinct aspects of neural organization change or recover.

Our work extends mechanistic studies of hormonal action beyond traditional work in receptor-rich hypothalamic circuits^6,7,16,37,73,74^. Social and reproductive behaviors are often studied through hormonally regulated innate actions such as sexual receptivity^6,7,16,22^, yet these behaviors also require continuous integration of sensory information, internal state, context, and competing drives and behavioral options. Our recordings during the social-choice paradigm captures this ongoing decision process and reveals that estrogen regulates not only which social option is preferred, but also how persistently that choice is maintained. At the neural level, reduced neural state flicker and longer intrinsic timescales provide a correlate of this increased persistence. Because mPFC contributes broadly to context-dependent choice, evidence accumulation, and flexible behavioral control, similar principles may extend beyond social behavior to other forms of decision-making influenced by internal physiological state.

The computational features identified here may also represent shared principles of hormone-dependent neural remodeling across brain regions and sexes. Across brain regions, previous studies have reported hormone-dependent changes that parallel individual components of this framework, including changes in the persistence of hypothalamic population dynamics across the estrous cycle^22,75^, increased hippocampal spine density and representational stability during high-estrogen states^2^, and reorganization of hypothalamic parenting nodes across pregnancy^5,29,76^. Human longitudinal imaging studies likewise indicate widespread changes in functional connectivity and network organization across menstrual-cycle^3^, pregnancy^18,32^, and other hormone-associated transitions including puberty and menopause^19^. Similar longitudinal dense-sampling of a male human brain^31^ revealed connectivity changes that closely track testosterone fluctuations, which occur on a faster, circadian timescale. Together with our findings, these observations raise the possibility that hormonal state can repeatedly tune neural systems along dimensions of stability, flexibility, and representational organization across circuits and physiological transitions. Determining which elements of this framework generalize across puberty, pregnancy, postpartum, perimenopause, and across sexes and species will be important for establishing what set of common computational principles govern endocrine remodeling of the brain.

Our study motivates future perturbation studies to test for robust attractor states^77^, and directly manipulate local recurrent architecture^78^ to test how specific connectivity motifs contribute to coding and persistence. Combining ERα knockdown with longitudinal recordings could further resolve the relative contributions of local mPFC remodeling and hormone-sensitive upstream inputs to these network-wide changes. A second opportunity is to improve temporal resolution of hormonal state read-out. Estrous staging provides limited temporal precision, whereas estradiol priming could reliably distinguish low- and high-estrogen conditions but still involve individual variability in hormone kinetics and return to baseline. Using recently developed optical reporters of estrogen-receptor-dependent transcription^1^ could provide continuous, local measurements of estrogenic signaling and reveal whether individual differences in transcriptional dynamics predict variation in connectivity and neural coding. Finally, naturally occurring hormonal cycles involve coordinated fluctuations in multiple hormones^4,29^. While the induced-cycle experiments isolate a major contribution of estrogen, an important first step in developing a computational framework for hormonal action, future studies can build on this framework to determine how progesterone^79^ and androgens^80^ shape the timing, magnitude, and reversibility of cortical reconfiguration across physiological states.

Hormonal action has classically been viewed as modulating specific receptor-expressing cells to gate particular behavioral outputs. Our findings support a broader view in which the endocrine state can shift an entire cortical network between persistent and flexible regimes, coordinating changes in connectivity, single-neuron coding, population dynamics, and behavior across slow timescales. This cross-scale framework may ultimately help provide biomarkers and interventions for maladaptive plasticity during profound hormonal transitions.

## Author Contributions

Conceptualization: M.M.A, A.L.F. Data curation: M.M.A, A.L. Formal analysis: M.M.A, D.T. Funding acquisition: M.M.A., A.L.F. Investigation: M.M.A, A.L, M.C, L.S., F.D.R. Methodology: M.M.A, D.T., S.O, F.D.R., A.L. Resources: A.L.F. Supervision: A.L.F. Writing - original draft: M.M.A. Writing - review & editing: M.M.A, A.L.F.

## Data availability

Data are available from the corresponding author upon reasonable request. All data required to reproduce the analyses in this manuscript will be made available on FigShare following publication of this manuscript.

## Code availability

Relevant code is available on GitHub repository: https://github.com/FalknerLab/Estrogen-mPFC-SocialChoice

## Acknowledgments

We thank E. M. Guthman for training on gonadectomy; T. Engel, C. Zimmerman, T. Z. Luo and members of the Falkner laboratory for useful discussions; J. Iravedra-Garcia, D. Allen and B. Mimica for feedback on the manuscript; staff at the PNI Viral Core Facility for reagents and the PNI Brain Registration and Histology Core Facility for assistance with histology; S. Kumar for mouse cartoon illustration. Funding was from Jane Coffin Childs Memorial Fund (M.M.A), Charles H. Revson Foundation (M.M.A), NIH DP2MH126375 (A.L.F.), NIH R01MH126035 (A.L.F.), NYSCF (A.L.F.), Simons Foundation (SCGB) (A.L.F.), Klingenstein Foundation (A.L.F.), and Alfred P. Sloan Fellowship (A.L.F.), McKnight Foundation (A.L.F), Allen Institute (A.L.F), One Mind Foundation (A.L.F.). A.L.F. is a New York Stem Cell Foundation Robertson Investigator.

## Declaration of Interests

The authors declare no competing interests

## FIGURES

**Extended Data Fig. 1.**
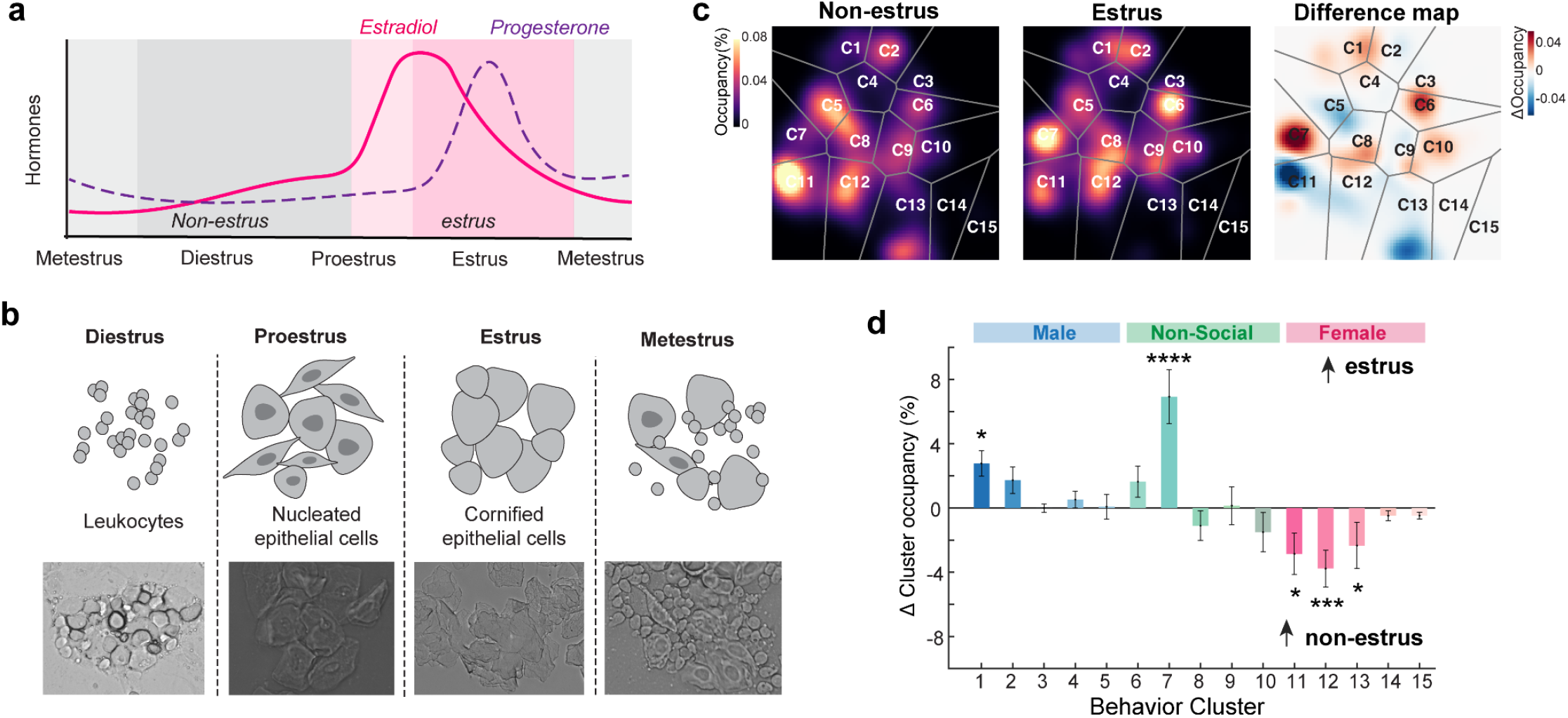
Estrous staging and changes in behavioral maps and finer-cluster occupancy across the endogenous cycle. **a,** Schematic of the endogenous hormonal cycle in naturally cycling mice. **b,** Schematic (top) and representative vaginal cytology images (bottom) for each phase of the estrous cycle. **c,** Occupancy density maps in UMAP space averaged across non-estrus and estrus sessions for an example mouse, with the difference (estrus − non-estrus) shown at right. Color scale, percentage of session time per bin (Δ percentage points for the difference map); grey lines, cluster boundaries. **d,** Change in behavioral-cluster occupancy during estrus. Bars denote mean Δ occupancy during estrus relative to each mouse’s mean non-estrus occupancy, expressed as percentage points of session time. (n = 8 mice, 16 estrus sessions; linear mixed-effects model accounting for repeated sessions within mice, with FDR correction across clusters.)

**Extended Data Fig. 2.**
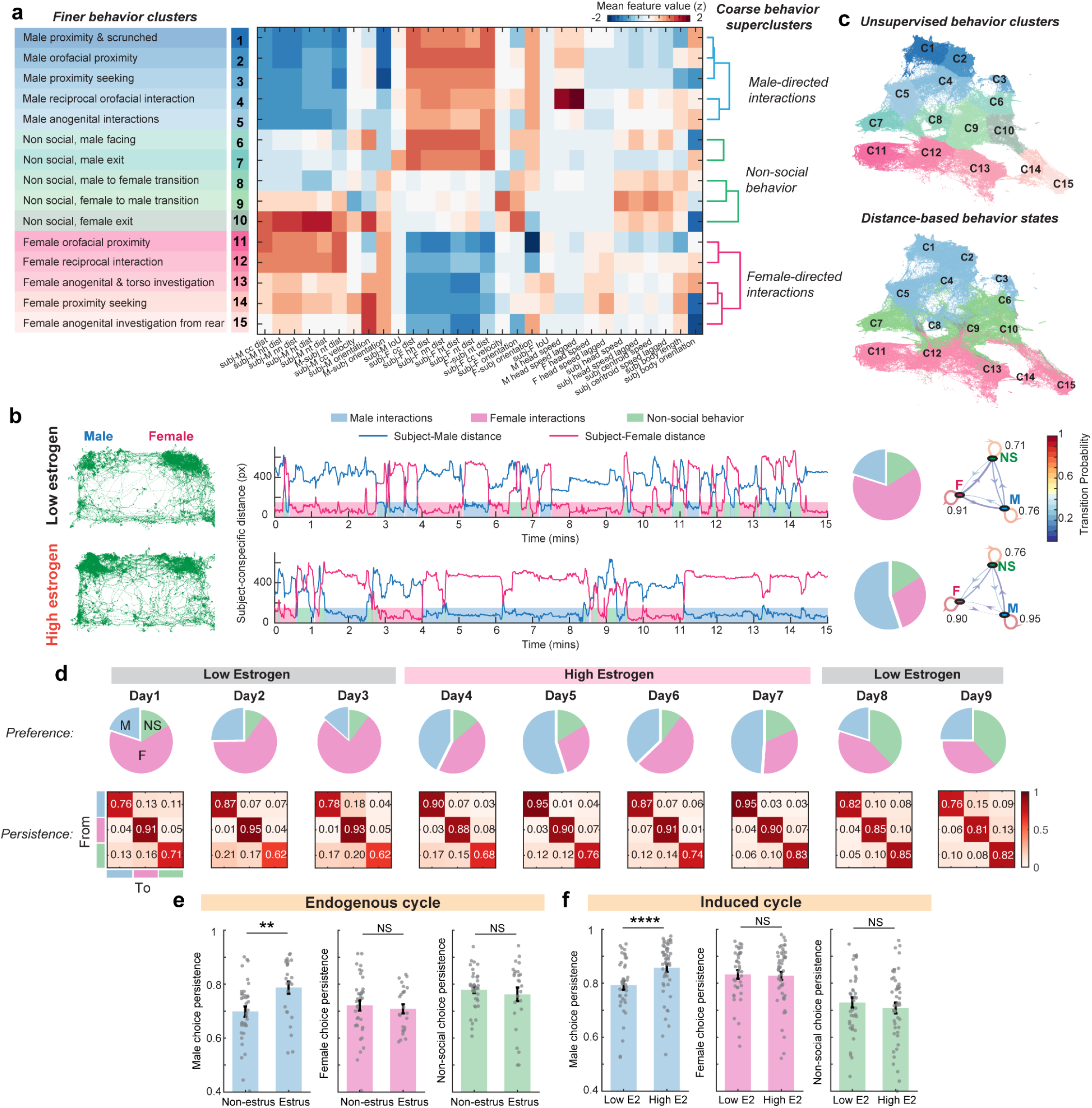
Unsupervised and distance-based characterization of behavioral states during the social-choice assay. **a,** Finer behavioral-cluster identities in UMAP space and mean z-scored values of SLEAP-derived behavioral features used for the embedding (left). Coarse behavioral superclusters obtained by hierarchical clustering of finer clusters based on normalized feature values, with corresponding dendrograms (right). subj., subject; M, Male conspecific; F, Female conspecific, dist., distance; c, centroid; h, head; n, nose; t, tail; IoU, intersection over union. **b,** Subject head position during example low E2 and high E2 sessions (left). Centroid-to-centroid distance between the subject and male and female conspecifics, with distance-based behavioral-state assignments shown below (middle). Pie charts show percentage of time in each behavioral state and transition graphs show probabilities of remaining in or transitioning between states (right). **c,** Correspondence between unsupervised and distance-based behavioral-state characterization. UMAP embedding color-coded by coarse behavioral superclusters and finer clusters (top), and by behavioral states obtained from the distance-based analysis in b (bottom). **d,** Pie charts used to quantify social preference and transition matrices used to quantify choice persistence across consecutive low E2 and high E2 recording days in an example mouse. **e,** Mean persistence during male-directed, female-directed and non-social behaviors across hormonal states in endogenously (left) and induced cycling mice (right). (Endogenous cycle: n = 13 mice, 37/25 non-estrus/estrus sessions; Induced cycle: n = 13 mice, 39/50 low/high E2 sessions; linear mixed-effects models accounting for repeated sessions within mice, with FDR correction.)

**Extended Data Fig. 3.**
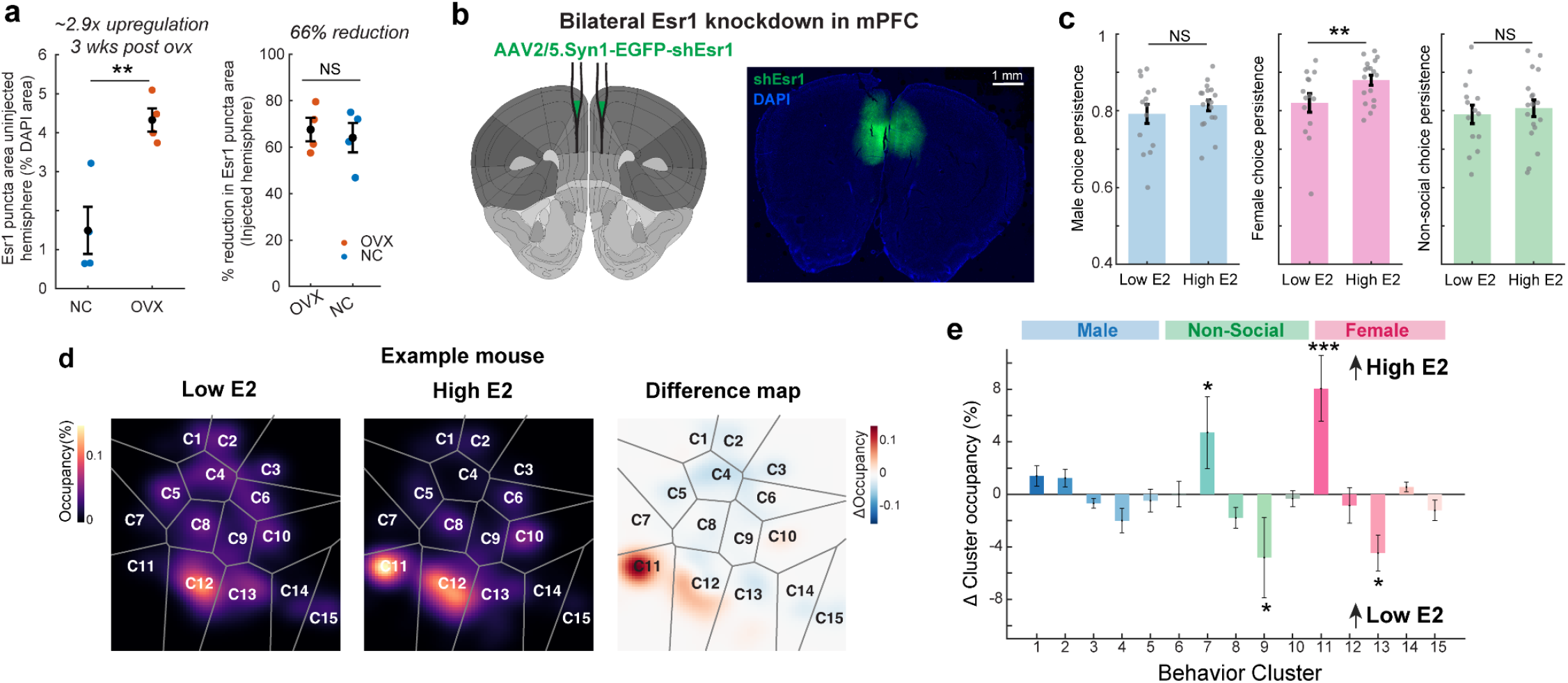
mPFC estrogen receptors are required for regulation of social choice. **a,** Esr1 puncta area, expressed as a percentage of DAPI area in the uninjected hemisphere, in naturally cycling and OVX mice 3 weeks after ovariectomy (left). Reduction in Esr1 puncta area relative to the uninjected control hemisphere (right); each point denotes one mPFC section. (n = 2 mice, 4 sections per mouse; two-sided two-sample t-test.) **b,** Bilateral mPFC injection for viral-mediated Esr1 knockdown using short-hairpin RNA (shEsr1). **c,** Mean persistence during male-directed, female-directed and non-social behaviors across low E2 and high E2 sessions in OVX mice with mPFC Esr1 knockdown. (n = 7 mice, 14/18 low/high E2 sessions; linear mixed-effects models accounting for repeated sessions within mice, with FDR correction.) **d,** Occupancy density maps in UMAP space averaged across low E2 and high E2 sessions for an example mouse, with the difference (high − low) shown at right. Color scale, percentage of session time per bin (Δ percentage points for the difference map); grey lines, cluster boundaries. **e,** Change in behavioral-cluster occupancy during high E2. Bars denote mean Δ occupancy during high E2 relative to each mouse’s mean low E2 occupancy, expressed as percentage points of session time. (n = 7 mice, 18 high E2 sessions; linear mixed-effects model accounting for repeated sessions within mice, with FDR correction across clusters.)

**Extended Data Fig. 4.**
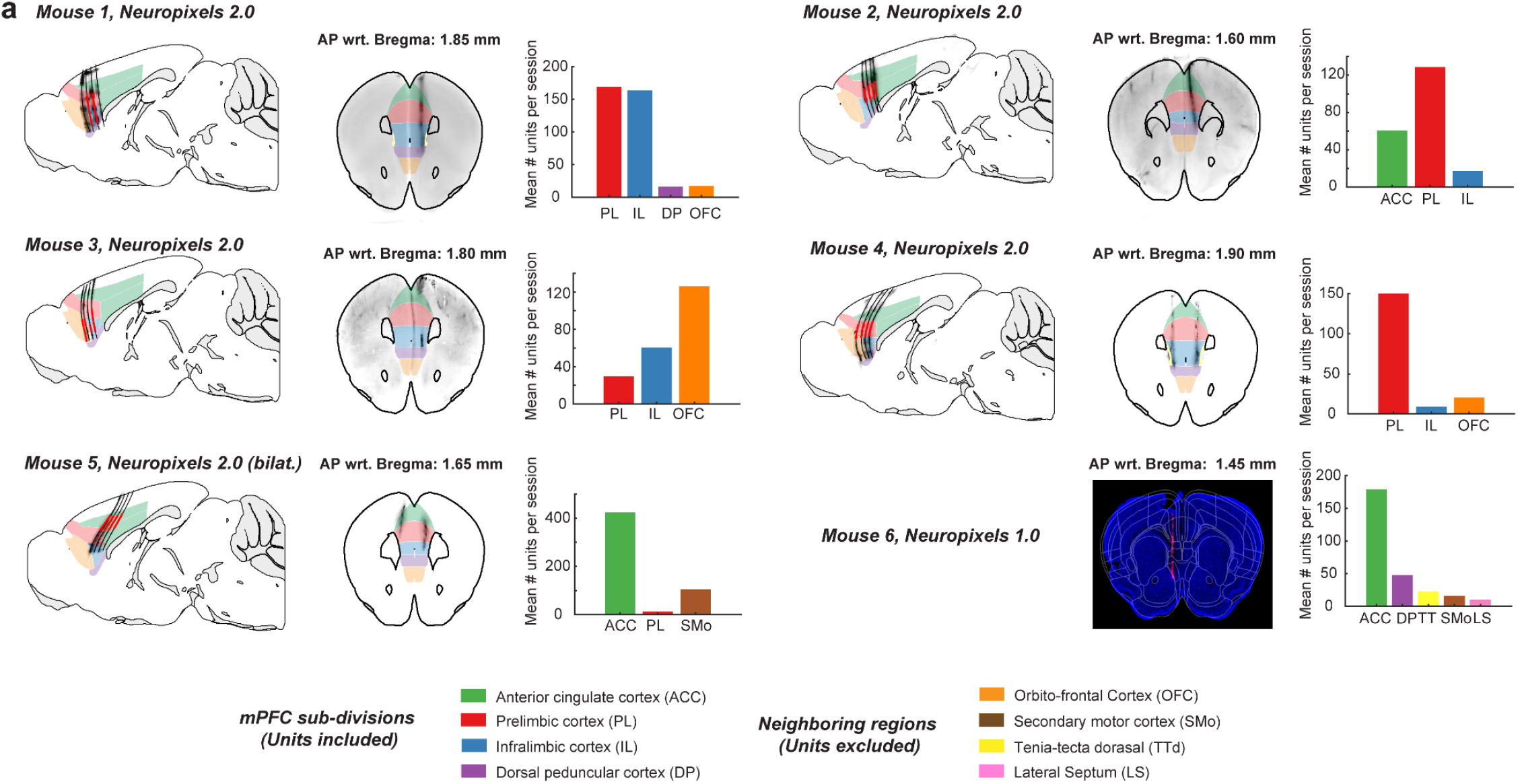
Chronic Neuropixels probe reconstructions and single-unit yield across mPFC subdivisions. **a,** Neuropixels 2.0 four-shank trajectories from mice 1-5 reconstructed using light-sheet microscopy after brain clearing, and a Neuropixels 1.0 single-shank trajectory from mouse 6 reconstructed using epifluorescence imaging of brain sections, registered to the Allen Common Coordinate Framework. Anatomical distribution and yield of isolated single units across mPFC subdivisions are shown alongside. Units recorded from neighboring regions are shown but were excluded from analyses.

**Extended Data Fig. 5.**
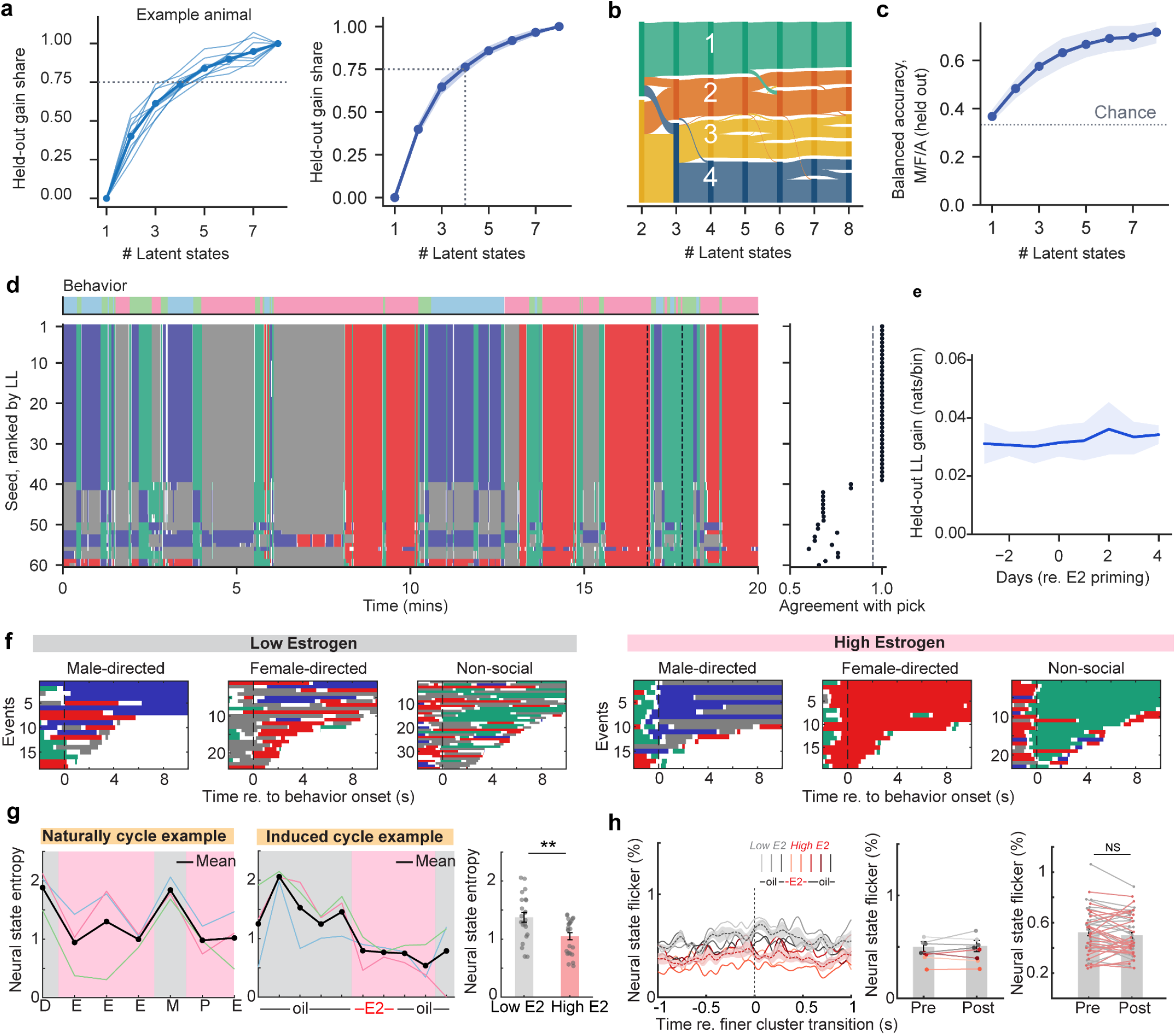
Hidden Markov Model states selection, model fit, neural state entropy and flicker a, Held-out gain share versus number of HMM states. HMMs were fitted independently with K = 1–8 states under five-fold cross-validation across days. The gain share is defined as the held-out log-likelihood gained over the one-state model, as a fraction of the gain at K = 8 (ℓ(k) − ℓ(1)) / (ℓ(8) − ℓ(1)). Dotted lines indicate where the pooled curve reaches 0.8 (K = 4.4) and 0.9 (K = 5.7) (Mean, shaded area 95% CI; n = 56 days, 6 animals.) The right panel shows the held-out gain share in only one example animal. **b,** HMM state lineage from K = 2–8. One example session’s HMM states (Animal 4, first E2 day), from fits at K = 2–8 with the same fold held out. Boxes are states (height = time occupied); bands connect states at K - 1 and K that share time bins. Colors and numbers identify the K = 4 states; at every other K, each box takes the color of the K = 4 state it overlaps most. **c,** Held-out behavior decoding from HMM states. For each K=1–8, each HMM state was assigned the behavior label with the highest P(behavior | state) in the training folds. Behavior was then predicted on the held-out fold from each bin’s decoded state and scored as balanced accuracy (mean recall over male, female and asocial); dotted line, chance (1/3). (Mean, shaded area 95% CI; n = 56 days, 6 animals.) **d,** Example HMM (K = 4) refits across random initializations. One example session (Animal 4, first E2 day), fitted from 60 random initializations. Each row is one initialization, ranked by training log-likelihood, where row 1 is selected as the best model (left). The fraction of time bins assigned the same state as the best (right; dashed line: 0.95). **e,** Held-out fit of the four-state HMM across days. Held-out log-likelihood gain over the one-state model per day (five-fold cross-validation, 60 random initializations). Days are aligned to the first E2 day (day 0) (Mean, shaded area 95% CI) **f,** Neural state rasters aligned to the onset of each behavioral state for an example low E2 (left) and high E2 (right) session. Behavioral events are sorted by length of the epoch. **g,** Neural state entropy within each behavioral state and averaged across states (black) over consecutive days in an example naturally cycling (left) and induced cycling mouse (middle). Mean neural state entropy across low E2 and high E2 sessions (right). (n = 6 mice, 22/24 low/high E2 sessions.) **h,** Neural state flicker aligned to finer behavioral-cluster transitions, expressed as a percentage of the total number of transitions, in an example mouse (left). Mean flicker in 1-s window before and after finer-cluster transitions (middle) and summary across sessions (right). (n = 6 mice, 22/23 low/high E2 sessions.)

**Extended Data Fig. 6.**
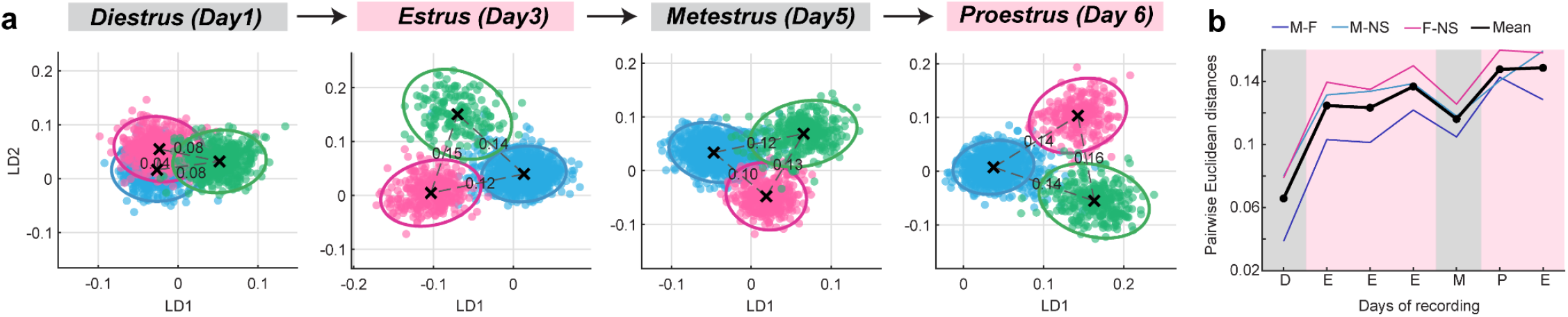
Estrogen increases neural-state persistence and population-level discriminability of behavioral states across the endogenous cycle. **a,** Population activity projected into linear discriminant (LD) space during example sessions from different estrous stages. Each point represents population activity in one time bin, colored by the corresponding male-directed (blue), female-directed (pink) or non-social (green) behavioral state. Pairwise Euclidean distances are shown between class centroids. **b,** Euclidean distances between behavioral-state representations for each pairwise comparison and the mean across comparisons (black) over consecutive estrous-cycle days in an example mouse.

**Extended Data Fig. 7.**
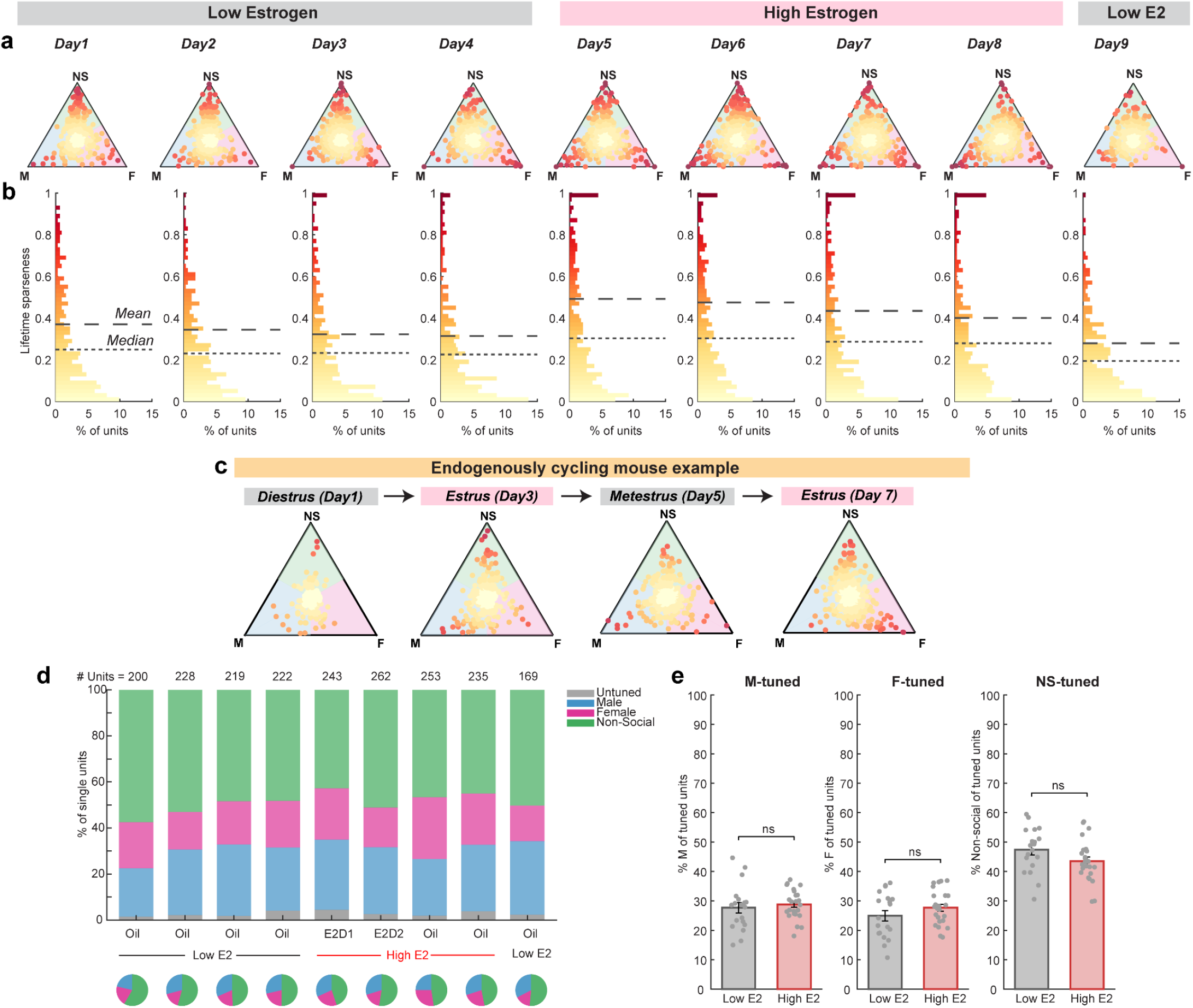
Estrogen increases single-unit tuning selectivity without altering tuning-class composition. **a,b,** Single-units from each session projected onto the tuning triangle (a); each point denotes one unit and is colored by lifetime sparseness. Corresponding lifetime-sparseness distributions are shown below (b). **c,** Single units recorded during example sessions from different estrous stages projected onto the tuning triangle for an example naturally cycling mouse. **d,** Stacked bar plots and pie charts showing the percentage of units tuned to each behavioral state across consecutive recording days; total number of units recorded each day is shown above. **e,** Percentage of units tuned to each behavioral state across low E2 and high E2 sessions. (n = 6 mice, 19/24 low/high E2 sessions; linear mixed-effects models accounting for repeated sessions within mice, with FDR correction.)

**Extended Data Fig. 8.**
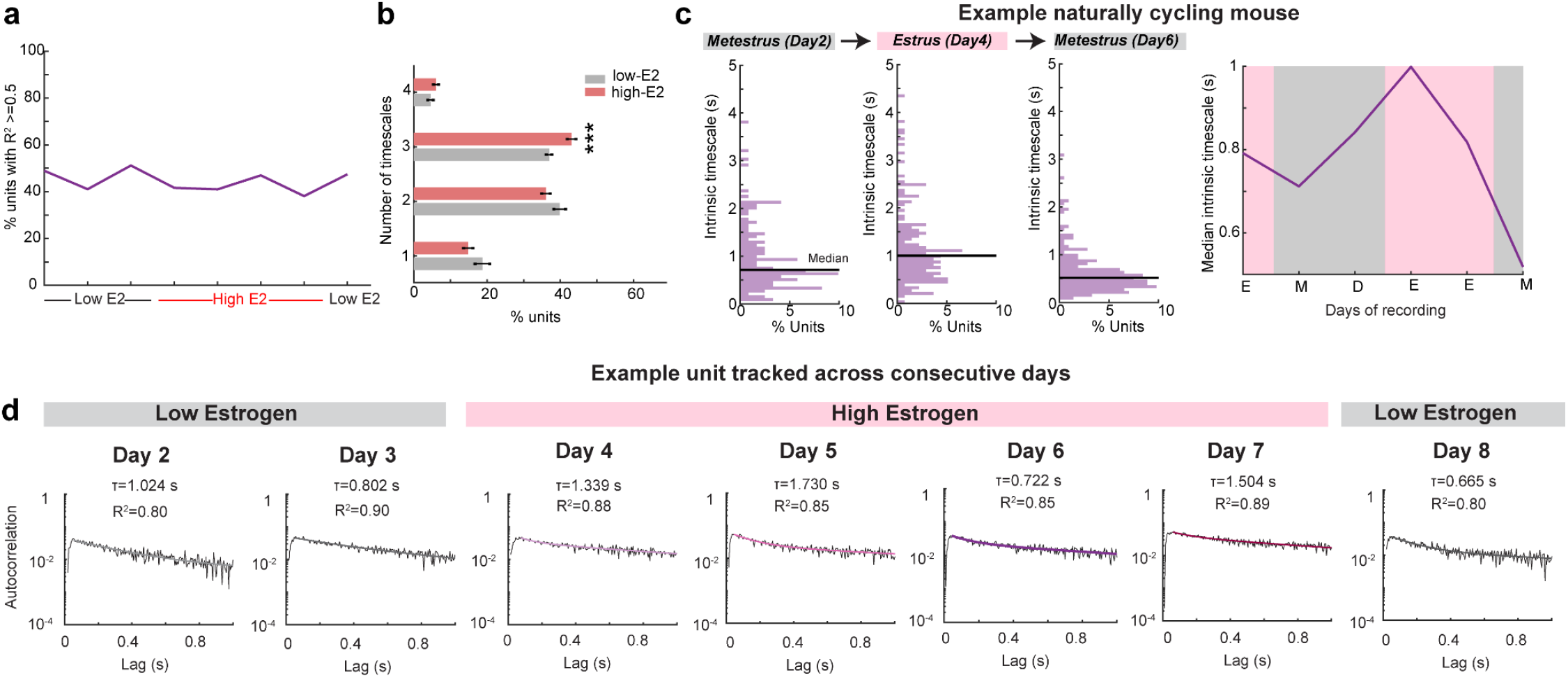
Estrogen increases intrinsic timescales in endogenous cycling mice and for tracked units. **a,** Percentage of units included in intrinsic-timescale analyses across consecutive sessions based on goodness of fit of the mixture-of-exponentials model to the autocorrelation function. **b,** Percentage of units best fit by models containing one to four timescales across low E2 and high E2 sessions. (n = 6 mice, 19/21 low/high E2 sessions; linear mixed-effects models accounting for repeated sessions within mice, with FDR correction.) **d,** Distribution of intrinsic timescales during example sessions from an example naturally cycling mouse (left), and the median intrinsic timescale over consecutive estrous-cycle days (right). **e,** Autocorrelation data (grey) and corresponding mixture-of-exponentials fits for an example single unit tracked across seven consecutive low E2 and high E2 days; normalized autocorrelation functions for the same unit are shown in Fig. 4g.

**Extended Data Fig. 9.**
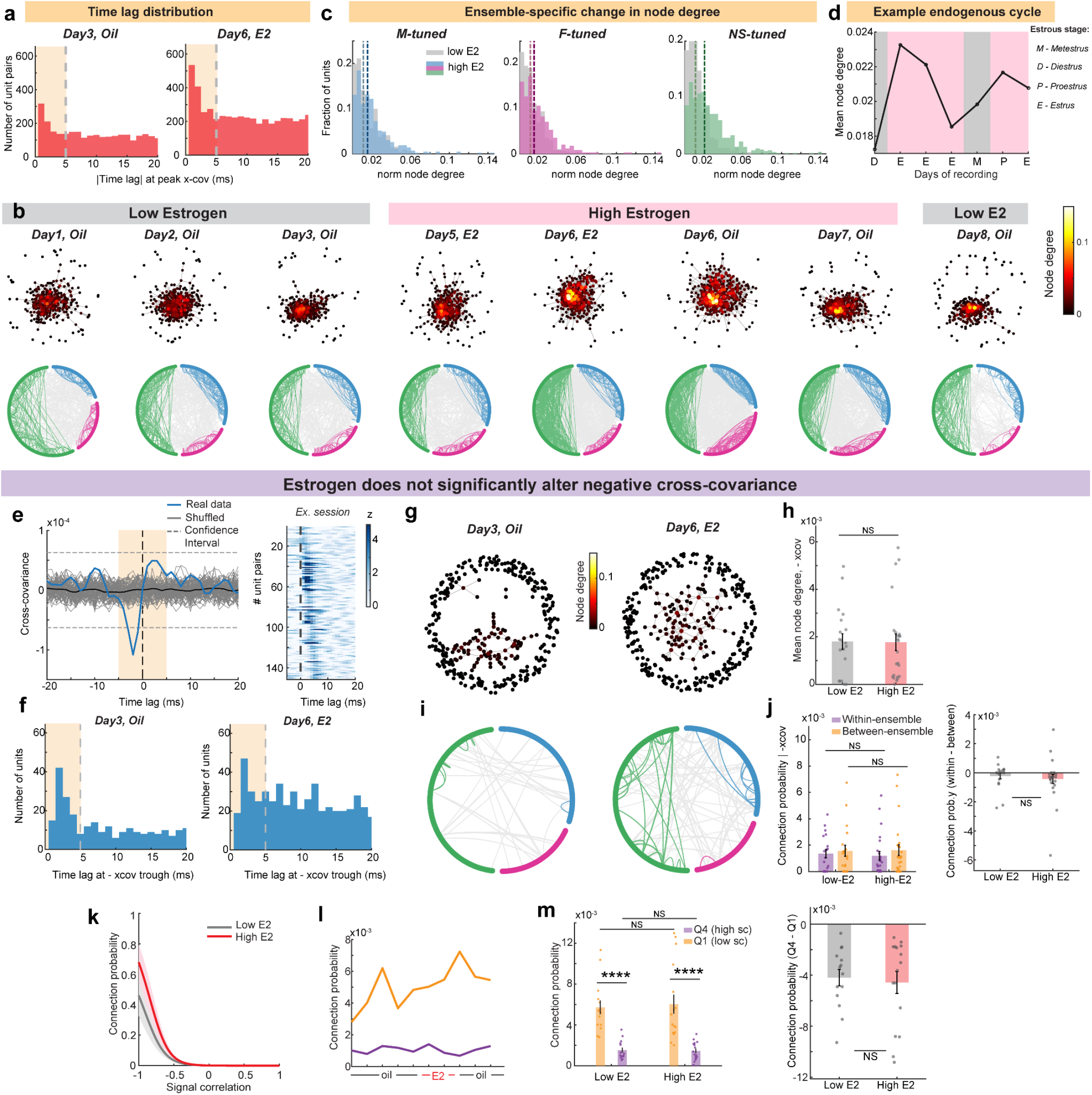
Estrogen strengthens excitatory resting-state connectivity in mPFC without altering negative cross-covariance. **a,** Distributions of absolute time lags at which spike cross-covariance functions peak during example low E2 and high E2 sessions. **b,** Network graphs depicting resting-state functional connectivity across consecutive low E2 and high E2 recording days in an example mouse (top). Nodes denote single units, edges denote significant cross-covariance and node color indicates normalized node degree. Corresponding chord plots are shown below; dots denote units grouped by tuning, arcs denote significant connections, within-ensemble connections are shown in the corresponding tuning-group color and between-ensemble connections in grey. **c,** Node-degree distributions for units tuned to each behavioral state during low E2 and high E2 sessions in an example mouse. **d,** Mean normalized node degree over consecutive recording days in an example naturally cycling mouse. **e,** Example negative spike-time cross-covariance for a pair of single units during spontaneous activity (left). Significant inhibitory connections were defined by cross-covariance below the confidence interval at a non-zero |lag| ≤ 5 ms. Normalized negative cross-covariance for all significantly connected unit pairs from an example session is shown at right. **f,** Distributions of absolute time lags at which negative spike cross-covariance functions show a significant trough during example low E2 and high E2 sessions. **g,** Network graphs depicting resting-state inhibitory functional connectivity during example low E2 and high E2 sessions (left). Nodes denote single units, edges denote significant negative cross-covariance and node color indicates normalized node degree. **h,** Mean normalized node degree for inhibitory connections across low E2 and high E2 sessions (right). (n = 6 mice, 19/24 low/high E2 sessions; linear mixed-effects model accounting for repeated sessions within mice.) **i,** Chord plots for example low E2 and high E2 sessions (left). Dots denote single units grouped by tuning, arcs denote significant inhibitory connections, within-ensemble connections are shown in the corresponding tuning-group color and between-ensemble connections in grey. **j,** Inhibitory connection probability for within-ensemble and between-ensemble unit pairs across low E2 and high E2 sessions (right). (n = 6 mice, 19/24 low/high E2 sessions; linear mixed-effects models accounting for repeated sessions within mice, with FDR correction.) **k,** Connection probability as a function of signal correlation (sc) for unit pairs with significant positive (top) and negative (bottom) cross-covariance, fit using logistic regression for an example mouse (left). Curves show mean fits across sessions and shaded regions indicate s.e.m. **l-m,** Mean connection probability for unit pairs in the lowest (Q1; low sc) and highest (Q4; high sc) quartiles across consecutive recording days (l) and across low E2 and high E2 sessions (m). (negative cross-covariance: n = 4 mice, 14/16 low/high E2 sessions; linear mixed-effects models accounting for repeated sessions within mice, with FDR correction.)

**Extended Data Fig. 10.**
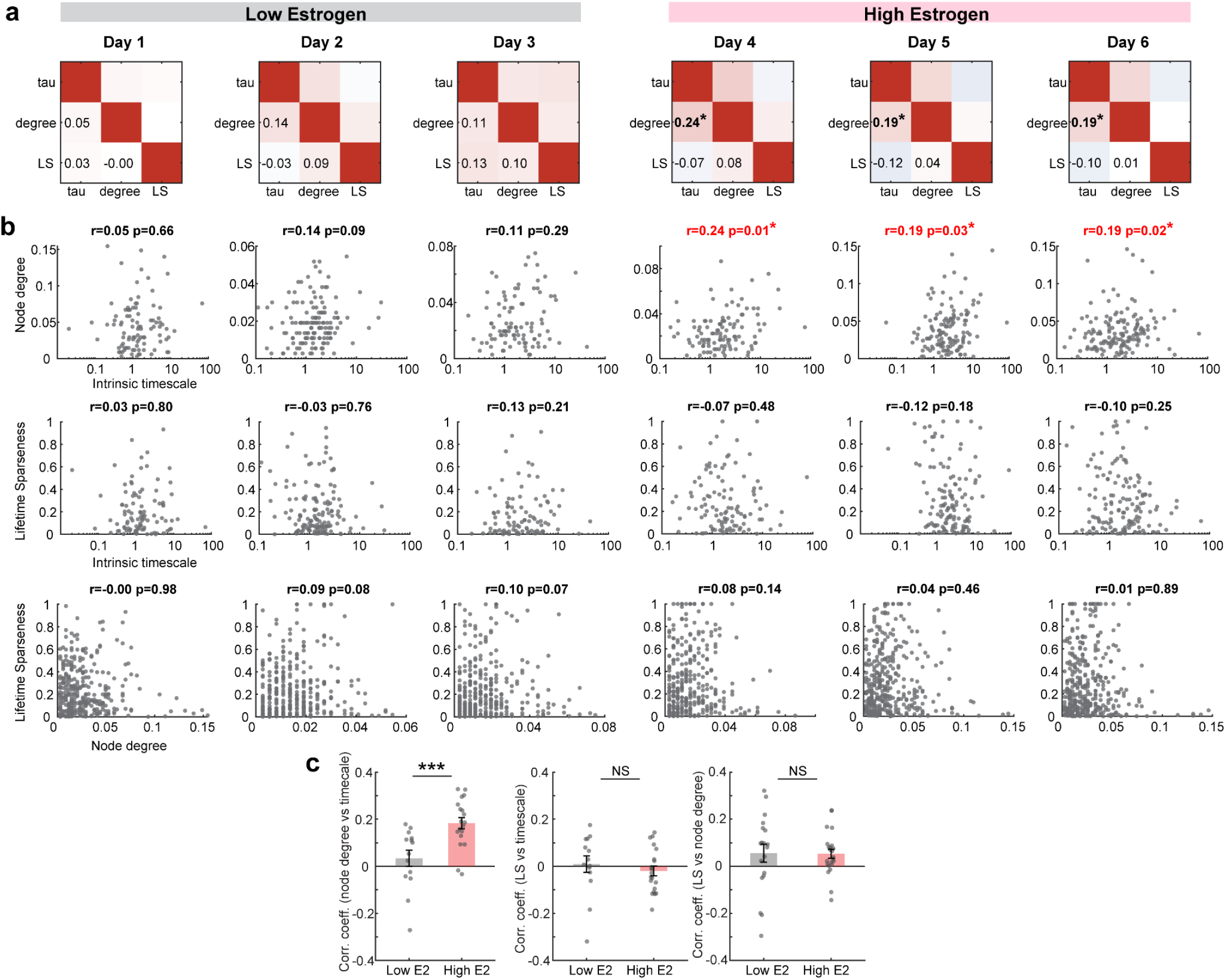
Relationships between intrinsic timescales, node degree and tuning selectivity. a, Correlation coefficients between intrinsic timescale, normalized node degree and tuning selectivity for single units recorded during each low E2 and high E2 session for example mouse. **b,** Session-wise correlation between intrinsic timescale and normalized node degree (top), intrinsic timescale and lifetime sparseness (middle), normalized node degree and lifetime sparseness (bottom). *r* denotes Spearman’s correlation coefficient; session-wise coefficients are shown as uncorrected descriptive statistics. **c,** Mean correlation coefficients across low E2 and high E2 sessions for all three comparisons (n = 5 mice, 14/19 low-E2/high-E2 sessions for intrinsic timescale versus node degree; 5 mice, 14/20 low-E2/high-E2 sessions for intrinsic timescale versus lifetime sparseness; n = 6 mice, 19/23 low-E2/high-E2 sessions for node degree versus lifetime sparseness; linear mixed-effects model accounting for repeated sessions within mice.)

**Supplementary Table 1.**

Information outlined in each section contains the figure and panel number (columns 1-2), a description of the measurement and comparisons (columns 3-5), sample sizes (column 6), statistical test employed (column 7), outcome with relevant statistics and p-values (columns 8-9) and post-hoc comparisons or adjustments (columns 10-13). P-values in the manuscript are reported as follows: *p < 0.05, **p<0.01, ***p<0.001, ****p<0.0001, N.S. if not significant.

## METHODS

### Mice

All animal procedures were approved by the Princeton University Institutional Animal Care and Use Committee and were in accordance with National Institutes of Health Standards. Adult female wildtype CD1 strain mice (Charles River Strain Code 022, Massachusetts, USA) aged 10-16 weeks at the start of experiments were used for all studies. Mice were typically group-housed with 2-4 same-sex mice per cage at 21-26 °C and 30-70% humidity in a 12-hour reversed dark-light cycle. Mice that were implanted with Neuropixels were single-housed after the implantation surgery to preserve the probe and headstage. Experiments occurred exclusively during the subjective active (dark) phase. Food and water were provided ad libitum.

All experimental subjects were sexually naive at the start of experiments. For mice used as social targets (partners), age matched sexually naive wildtype CD1 strain males and females were used. All subject mice were naive to partner mice prior to interaction. All animals were bred within the Princeton University animal facilities or purchased directly from Charles River Laboratories.

#### Estrous staging in naturally cycling mice

The estrous cycle was monitored on experimental days for all naturally cycling female subjects and partner mice. Vaginal epithelial cells were collected using a cotton swab with care taken to avoid deep penetration that might induce pseudopregnancy. Vaginal cytology was subsequently imaged under a brightfield microscope, and mice were staged based on the observed cell types. Mice with nucleated epithelial cells were assigned to proestrus, mice with cornified epithelial cells were assigned to estrus, and mice with leukocytes were assigned to metestrus or diestrus depending on proportion of leukocytes and cornified/nucleated cells. For comparisons between estrus and non-estrus, data from proestrus and estrus days were combined into the former category and data from metestrus and diesterus days were combined into the latter category.

#### Hormonal priming in ovariectomized mice

Ovariectomized mice recovered for 2-3 weeks before beginning the hormonal priming regimen. Before initiating estradiol priming, low-estrogen baseline was first established by performing daily subcutaneous injections of 25 uL of sesame oil (Sigma-Aldrich, CAS #8008-74-0) to the mice over 3 to 5 days. During estradiol (17β-Estradiol, Sigma-Aldrich, CAS #50-28-2) replacement,10ug on Day 1 and 5ug on Day 2 was administered as subcutaneous injections of the estradiol solution in sesame oil (2mg/5ml). These levels of estradiol have been previously shown to be comparable to physiological proestrus peak levels^6,14^. Behavioral testing or electrophysiological recordings were all conducted approximately 4-5 hrs after the injection.

### Behavior

The recording chamber consisted of a custom box (20” x 10”, 17”) made of acrylic boards (TAP Plastics Acrylic Sheets P95 Matte Finish - Matte Black 1/4” thick) laser cut to the specific dimensions. The front wall is transparent to allow for video recordings from the front as well. Subjects have a 10 min “resting-state” period in the chamber to allow for habituation and recording of task- and stimulus-free spontaneous neural activity each day prior to the start of the social choice assay.

#### Social choice assay

Female subject mice undergo a 15-20 minute social choice assay each day of recording where they are free to choose and interact either with a male or a female conspecific tethered to two side-by-side corners of the recording chamber. The conspecifics are age-matched, sexually naive and novel to the subject prior to the first day of behavior. The same male and female conspecifics are tethered on the same side of the arena for the rest of the recording sessions for a given subject, and the sides are counterbalanced across different subjects. The conspecifics were constrained in custom full body jackets (made from ThorLabs BK5 nylon fabric, cut on an Epilog Mini 24 30W laser) and tethered to the back wall of the recording chamber using a custom set-up with fishing line (Piscifun Onyx Braided Fishing Line, 150lb), jewelry clasps (uxcell 12mm Round Stainless Steel Spring Ring Clasps) and magnets (KeySmart MagConnect Magnetic Key Holder). Jackets were cut with holes so that the subject could have substantial access to the conspecific’s body surface during the interactions, and yet the conspecifics were comfortably constrained. This allowed the subjects to investigate, sniff and groom the anogenital, torso and orofacial body parts of the conspecifics. The mice are briefly anesthetized during jacketing using isoflurane for ∼5 mins and are allowed to recover and are fully awake prior to the start of recordings. They are also well habituated to being jacketed prior to the first day of behavior.

#### Video acquisition

Behavioral sessions were recorded using up to eight cameras (BFS-U3-13Y3m-C1.3 Mono, FLIR Systems) acquired with campy, a python package for synchronized video acquisition and real-time data compression (https://github.com/ksseverson57/campy). The cameras are controlled with a Raspberry Pi and videos are recorded at 60 frames/sec.

### Surgery

For all procedures, mice were placed in an induction chamber under 3-5% isoflurane for several minutes to induce anesthesia and then injected with ketoprofen subcutaneously to provide analgesia. They were then placed on a stereotaxic frame (Kopf Instruments) and anesthesia was maintained with 1–2% isoflurane flowing continuously through a nose cone over the bite bar.

#### Ovariectomy

Ovariectomy, involving surgical removal of ovaries was performed using a lateral flank approach. Mice were brought to surgical levels of anesthesia as described above. In addition to ketoprofen, extended-release Buprenorphine was administered. Single incisions were made along the left and right flanks of the mouse to cut through the skin, followed by the internal muscle layer. The fat pad surrounding the ovaries was gently pulled to exteriorize the ovaries from the body cavity, and the connection between the ovary and its corresponding uterine horn was ligated with absorbable suture (Coated Vicryl, Ethicon, Johnson & Johnson, New Jersey, USA). A cut was then made to remove each ovary. The internal muscle layer was then sutured using absorbable sutures, and the skin incision was closed with surgical wound clips. The clips were removed following healing 10 days after surgery. All mice recovered for 2-3 weeks prior to the initiation of post-ovariectomy experiments.

#### Chronic Neuropixels implantation

After verifying surgical levels of anesthesia, mice were secured in ear bars, shaved and cleaned with ethanol and betadine. A midline incision was then made with a scalpel, skin on the top of the skull was removed and the skull was cleared of all overlying tissue. The craniotomies for the ground and reference were drilled distant from the brain region to be recorded and the skull screws were inserted and cemented after verifying the impedance between the ground and reference screws is less than 3 kΩ and making sure there is a robust electrical connection with brain tissue. The skulls were leveled using the confluence of the midline and rostral rhinal vein (anterior vein that sits on the dorsal surface of the mouse brain between the olfactory bulb and prefrontal cortices) as the anterior position and lambda as the posterior position. Craniotomy was then made at the implantation site of the probe centered at stereotaxic coordinates, +1.8 mm AP, 0.45 mm ML relative to Bregma. Just before insertion, the probe was coated in lipophilic dye (DiI in 100% ethanol) to enable histological reconstruction of the probe locations. The silver wires soldered to the ground and external reference pads on each probe flex cable were connected to the ground and reference skull screws. Probes were manually lowered for the initial penetration and then lowered automatically at a rate of 2 microns per second to the final depth of 2.7 mm DV relative to the brain surface. Then we used dental cement and a viscous dental composite (Absolute Dentin, Parkell) to create a wall-like barrier surrounding the probe while leaving the probe shank exposed to air. Dentin was applied at low viscosity to fill all gaps and secure the entire implant assembly to the Metabond-coated skull. The ground wires were also covered by the dentin to prevent them being disturbed after the surgery, and the entire assembly is wrapped in electrical tape for noise shielding.

#### Viral injections

For Esr1 knock-down experiments, we injected AAV2/5-Syn1-EGFP-shEsr1 (titre, 3.3 × 10^13^ genome copies (GC) per ml; manufacturer, Princeton Neuroscience Institute (PNI) Viral Core Facility) into mPFC bilaterally for testing necessity of ERɑ for behavior and unilaterally for RNAscope-FISH validation. Mice were anesthetized, secured in ear bars and skulls were leveled as described above and craniotomy was made at coordinates +1.8 mm AP, 0.45 mm ML relative to Bregma. Pulled glass micropipettes were backfilled with the virus solution, lowered into the target site and virus was infused at 2 nl/s using a precision nanoliter injection system (Nanoject III, Drummond). For behavior experiments, we injected 250 nl at 1.6 mm DV relative to the brain surface in both hemispheres, and for RNAscope experiments, we unilateraly injected 250 nl each at two depths, 1.6 mm and 2.4 mm DV relative to the brain surface. Following surgery, we allowed 3-4 weeks for viral infection and transgene expression.

### Fluorescent In Situ Hybridization

Altered expression of Esr1 and eGFP was confirmed histologically with RNAscope Multiplex Fluorescent ISH V2. All probes are available via ACDBio. The probe for Esr1 (catalogue 478201) consisted of 20 ZZ pairs designed to target region 678-1723 of the *Mus musculus* estrogen receptor 1 gene, transcript variant 1 (accession number NM_007956.5). The probe for eGFP (catalogue 400281) consisted of 13 ZZ pairs designed to target region 628-1352 of the enhanced green fluorescent protein (accession number U55763.1).

Detailed methods for the RNAscope Multiplex *fISH* V2 can be found courtesy of ACD Bio (https://acdbio.com/sites/default/files/UM%20323100%20Multiplex%20Fluorescent%20v2_Rev%20D.pdf). Mice were deeply anesthetized with 5% anesthesia. Following anesthesia induction, animals were quickly perfused with nuclease-free 1X phosphate-buffered saline to clear blood from circulation. Brain tissue was then promptly extracted and immediately frozen in O.C.T compound on dry ice before storing at -80°C. Tissue was habituated to -20°C approximately 2-hours before sectioning. Tissue was then sectioned into 20µm sections, mounted directly to pre-chilled Superfrost Plus slides, and stored at -80°C until the time of *fISH*. We used the method for *fISH* following the linked guidelines from ACD Bio, as specified for Fresh-frozen tissue. In lieu of using 100% protease-IV, we diluted protease-IV to 10% with nuclease free-water to preserve tissue integrity.

### Chronic Neuropixels electrophysiology

#### Neuropixels probe assembly for chronic reusable implants

For recoverable and reusable chronic implantation, we used the “Apollo implant” design^81^. It primarily consists of 3D printed fixtures with three main components: a) probe mount that can hold either one or two probes b) skull connector which is fixed to the animal’s skull using a viscous dental composite (Absolute Dentin, Parkell) c) docking module which is used to attach the probe mount onto the skull connector. These are secured together using screws that could be removed at the end of the experiment to enable explantation and reuse. To avoid failures related to connecting/reconnecting the zero-insertion-force (ZIF) connector on the headstage during each experimental session, the headstage was left on the animal for the entire duration of the experiments. The total weight of the assembled fixture with probes was ∼3 g.

#### Recording channel configuration, data acquisition and synchronization

Chronic electrophysiological recordings in freely moving subjects were performed using Neuropixels 1.0 or 2.0 probes and commercially available Neuropixels acquisition hardware used in conjunction with PCI eXtensions for Instrumentation (PXI) hardware from National Instruments (including a NI PXIe-1071 chassis, NI PXIe-8381, as well as NI PXI-6341 I/O module along with NI BNC-2110 for recording analog and digital inputs).

We used SpikeGLX software (https://billkarsh.github.io/SpikeGLX/) to acquire the data. We use the skull screw penetrating the brain at a different location from the probe insertion site as our reference. The amplifier gain used during recording with the 1.0 probes was 500 and 2.0 probes was 100. The Neuropixels 1.0 single shank recording channels were chosen from the deepest 384 recording sites (‘bank 0’) such that they span the entire depth of mPFC. Neuropixels 2.0 electrode arrays have 384 selectable recording sites from 5210 total sites distributed across four 1 cm shanks^41^. Shanks are 250 um apart and each have two vertical columns of sites. Recording configurations for the Neuropixels 2.0 recordings were chosen across two or four shanks such that single-yield was maximized while spanning either a depth of 1440 um or 720 um respectively within mPFC. Recordings lasted 45 to 60 minutes. Mice were habituated to the recording chamber and conspecific interactions prior to implantation. After surgery, mice were given up to a week for recovery before the start of chronic recordings each day.

Experimental control signals (including camera triggers) were recorded simultaneously using the same system. Electrophysiological recordings were synchronized to camera strobe or behavioral task events by aligning to a common digital signal sync signal randomly flipping between high and low states (produced from an Arduino) accounting for both clock offset and drift.

#### Atlas alignment and probe track reconstruction

All probes were coated in CellTracker CM-DiI (Invitrogen, C7000) before implantation. After the conclusion of experiments, animals were euthanized and the brains cleared with the iDISCO+ protocol^82^. The cleared brains were then imaged on a light-sheet microscope (LaVision Ultramicroscope II) using LaVision BioTec ImSpector software (https://www.lavisionbiotec.com; v.7.0). Images were acquired using 488 nm (autofluorescence channel) and 561 nm (CM-DiI channel) excitation light with 10 µm between horizontal planes and 5.91 µm per pixel resolution. Atlas alignment then followed the IBL’s pipeline^83^ (https://github.com/int-brain-lab/iblapps/wiki). The autofluorescence volume was registered to the atlas using the Python package Brainreg (https://github.com/brainglobe/brainreg; v.0.4.0), and these transformations were then directly applied to the CM-DiI volume. Individual Neuropixels shank trajectories were then manually annotated in the atlas-registered CM-DiI volume using the Brainreg-segment Napari module (https://github.com/brainglobe/brainreg-segment; v.0.2.16). Every recording site was then localized to an Allen Common Coordinate Framework (CCF)^84^ coordinate. The alignment process was performed separately for each Neuropixels probe shank. For all analyses, we only included neurons from recording sites that were localized to the mPFC sub-divisions primarily spanning Anterior cingulate cortex (ACC), Prelimbic cortex (PL), Infralimbic cortex (IL), but a small proportion of units come from Dorsal peduncular cortex (DP), Tenia tecta dorsal (TTd) which is anatomically at the ventral limit of mPFC and shown to be functionally ‘mPFC-like’^85^.

#### Spike-sorting, single-unit quality control and curation

Raw data within the action potential band (soft high-pass filtered over 300 Hz) was denoised by common mode rejection (that is, subtracting the median across all channels), and spike-sorted using Kilosort 2.5^86^ (www.github.com/MouseLand/Kilosort2). Bombcell^87^ (https://github.com/Julie-Fabre/bombcell) was first used for semi-automated curation and quality assessment of the units, followed by manual verification of bombcell unit-classifications using custom code. High-quality single-units were defined with the following bombcell criteria:

1. Waveform has maximum of one trough and two peaks; Units are classified as non-somatic if the higher amplitude peak precedes the trough
2. Maximum waveform amplitude has a slope of at least −0.003 per micron across channels, to eliminate non-localized units
3. Waveform trough-to-peak duration is >100 μs and <1000 μs
4. Waveform has a minimum amplitude of 20 uV and amplitude during baseline is <30% the maximum amplitude
5. Less than 20% of spikes are missing due to drift as estimated by a gaussian fit of spike amplitudes
6. At least 300 spikes are recorded and a presence ratio of 0.7 (≥1 spike in ≥70% of 60-s bins)
7. Less than 10% refractory period violations (false positives) in a 2 ms refractory period window

Units failing to meet these criteria were classified as noise and discarded from all analyses. Units were classified as regular spiking versus fast spiking based on peak-trough interval exceeding a threshold of 0.45 ms.

#### Longitudinal tracking of single-units across days

To track the same neurons across multiple recording sessions, we used UnitMatch^42^ and DeepUnitMatch^43^ depending on the probe type used for each animal. The default pre-trained DeepUnitMatch model was trained exclusively on Neuropixels 2.0 four-shank probe recordings, and the model architecture assumes a specific four-column channel geometry. One animal in our dataset was recorded with a Neuropixels 1.0 single-shank probe, and UnitMatch was used for cross-session unit tracking. All remaining animals were recorded with Neuropixels 2.0 four-shank probes and were tracked using DeepUnitMatch with the default pre-trained model. Units were required to have a minimum of 1000 spikes per session and at least 100 spikes in each half of the recording to ensure within-session stability. Only units tracked across two or more sessions were retained.

UnitMatch identifies putative unit matches across recording sessions by computing pairwise similarity between all clusters detected in any two sessions. For each unit, 1000 raw waveforms were extracted per recording session and averaged to form a template. Six feature metrics were computed for every candidate pair: centroid distance, waveform similarity, centroid overlap, spatial decay similarity, amplitude similarity, and location trajectory similarity. Each metric was independently evaluated against distributions of known within-session (same-unit) and across-session (different-unit) pairs to estimate the probability that a given score arose from a true match versus a non-match. DeepUnitMatch replaces the hand-crafted feature metrics of UnitMatch with learned latent representations extracted by a deep convolutional neural network, enabling more robust matching in the presence of waveform variability due to probe drift or recording quality differences across sessions.

### Data exclusions

Animals were excluded if surgical probe placement was incorrect or if surgery-related issues compromised behavioral recordings. Single units were excluded based on Bombcell quality-control criteria and anatomical localization; units outside the mPFC were excluded from all neural analyses. Recording sessions were excluded when data acquisition or synchronization issues compromised data quality or behavioral recordings. Additional exclusions were applied where required by the criteria specific to individual analyses, and analysis-specific exclusion criteria and their rationale are described in the corresponding sections below.

### Behavior analysis

#### Pose tracking

Animal pose estimation was performed with SLEAP software (v1.2.8). To create a model, mice in a subset of frames from 485 videos were hand-labeled with a 15-node skeleton which labeled the following points: nose, head, neck, trunk, tail-trunk-interface, left ear, right ear, left forepaw, right forepaw, left hindpaw, right hindpaw, upper segment of tail, middle-segment of tail, lower segment of tail, and tail tip. All together, 19540 labeled frames were split into a training and validation set using an 80:20 split. Training of the SLEAP model was done using a top-down approach with customized parameters. We then used this model to run inference on all top-view behavior videos, whereby the model produced predictions that were then proof-read to produce final pose estimation.

#### Feature definition and preprocessing

For downstream behavior analysis, frame-to-frame pose data was converted into the following 30 behavioral features:

Social features (separately for the two partners):

1. Between centroid distance - Euclidean distance between the centroids of the subject and partners (centroid is defined as the mean of the 5 midline nodes: nose, head, neck, trunk and tail-trunk-interface)
2. Between nose distance - Euclidean distance between the nose of the subject and partners
3. Between head distance - Euclidean distance between the head of the subject and partners
4. Subject-to-conspecific head-to-tailbase distance - Euclidean distance between the head of the subject and the tailbase of the partner
5. Subject-to-conspecific nose-to-tailbase distance - Euclidean distance between the nose of the subject and the tailbase of the partner
6. Partner-to-subject nose-to-tailbase distance - Euclidean distance between the nose of the partner and the tailbase of the subject
7. Between centroid velocity - Instantaneous change in the between centroid distance (every 6 frames or 100ms)
8. Orientation of subject with respect to the partner - Angle between 2 vectors, one connecting tailbase and head of subject and second connecting head of subject and centroid of the partner
9. Orientation of partner with respect to the subject - Angle between 2 vectors, one connecting tailbase and head of partner and second connecting head of partner and centroid of the subject
10. Intersection over union of subject and partner (IoU): Overlap between subject and partner bounding boxes by dividing the area of their intersection by the area of their union. (bounding box is the box that bounds the nose-shoulder-haunch-tti (tail-trunk-interface)) Partner-specific features (separately for the two partners):
11. Partner head speed: Instantaneous change in distance of partner head position (every 6 frames or 100ms)
12. Partner head speed lagged at ± 3s: Sum across lags of partner head speed, computed within a ± 3s lag window. Subject-specific features:
13. Subject head speed: Instantaneous change in distance of subject’s head position (every 6 frames or 100ms)
14. Subject head speed lagged at ± 3s: Sum across lags of subject’s head speed, computed within a ± 3s lag window.
15. Subject centroid speed: Instantaneous change in distance of subject’s centroid position (every 6 frames or 100ms)
16. Subject centroid speed lagged at ± 3s: Sum across lags of subject’s centroid speed, computed within a ± 3s lag window.
17. Subject body length: Euclidean distance between the head and tailbase of the subject
18. Subject body orientation: Angle of the vector pointing from subject’s tailbase towards its head

Features for each session were processed by 1) removing outliers 3 standard deviations away from the session mean, 2) smoothing via Gaussian filter of width 0.25 s or 15 frames, 3) linear interpolation of missing values, and 4) z-scoring using the mean and standard deviation pooled from collating features across all sessions. The collated mean and standard deviation values for every feature were used for z-scoring in individual sessions to capture a wider variance in feature dynamics.

#### Unsupervised behavior analysis: Low-D UMAP embedding, finer and coarser cluster generation

To comprehensively characterize social behavior in an unbiased manner, we adapted previous unsupervised behavior classification strategies^37,45,46^ involving the reduction of high dimensional behavior features to a low-dimensional embedding and definition of behaviors as high-density clusters within this embedding. To do this, we embedded the 30 behavior features described above into a 2-dimensional embedding using UMAP (Uniform Manifold Approximation and Projection), a non-linear, graph-based dimensionality reduction technique that is based on manifold learning and topology. Briefly, it works by building a network based on how strongly points are connected to their nearest neighbors in high-dimensional space, and then finds a low-dimensional map that best preserves those connections.

To generate the UMAP embedding, an importance sampling approach was taken. This approach increased the likelihood of generating a final version of the embedding that also features rare or nuanced behaviors. Importance sampling altogether involved two rounds of UMAP reduction. In the first round, 98,400 video frames were uniformly sampled from 41 social choice assay videos across 4 mice. This total pool of feature frames was next embedded into a two-component UMAP manifold using parameters: number of neighbors = 15, minimum distance = 0.3, distance metric = euclidean. This was further parcellated using DBSCAN (Density-Based Spatial Clustering of Applications with Noise), a density-based clustering method that groups points based on how closely packed they are, using parameters: neighborhood radius, ε = 1.5, minimum number of points = 15, ε distance metric = euclidean, and this resulted in 15 clusters. For each point, if at least the minimum number of points are within ε radius, it is marked as a core point. Points reachable from a core point within epsilon are added to the same cluster, and points that aren’t reachable from any core point are labeled as noise.

As UMAP is non-parametric and does not provide a direct function to map new data points into the learned low-dimensional space, a separate model was implemented to learn the mapping from the original 30-dimensional feature space to the new 2D UMAP space, allowing the data from every video frame to be represented in UMAP space and thus fall into any of the 15 clusters previously defined. To enable consistent projection of all video frames into the same 2D UMAP embedding, we first computed UMAP on the training dataset and saved the resulting manifold mapping as a template file. This template captures the neighbor graph, local connectivity weights, and global optimization parameters that define the mapping from the original 30-dimensional feature space to the 2D embedding. Because the UMAP template only provides a projection into 2D space and does not assign cluster labels to new points, we trained a k-nearest neighbors (k-NN, k = 5) classifier using the 2D coordinates of the training data as inputs and the DBSCAN cluster labels as targets. The k-NN model then allows held-out frames to be projected into the UMAP space using the template and assigned to one of the pre-defined clusters based on the majority label of their nearest neighbors, ensuring that cluster assignments remain consistent without recalculating or altering the original clusters.

To generate the final UMAP embedding, over 20,000 frames were sampled from the 15 clusters generated previously. Specifically, 160 random frames from each cluster and for each recorded session were pooled together for a total of 96000 frames (160 frames x 15 clusters x 40 videos = 96,000). From these sampled frames, the 30-dimensional feature set was embedded into 2D UMAP space. Next, the full set of video frames for each mouse were mapped into this final UMAP manifold using the template and the k-NN model as described above. All behavior analyses involving behavior clusters are performed using this final UMAP manifold. Animals in which behavior was recorded using the older conspecific-tethering setup were excluded to ensure that a common master embedding was generated from behavior recorded using the same setup. Naturally cycling animals that remained in a single estrous-cycle phase were also excluded from UMAP analyses because occupancy comparisons required paired high- and low-estradiol conditions within each animal. All these animals were retained for distance-based behavioral analyses described below.

##### Comparison of behavior maps

To compare cluster occupancy between behavior maps, we used Jensen-Shannon (JS) divergence. JS divergence measures the similarity between two probability distributions (i.e., cluster occupancy) with more similar distributions approaching a JS divergence value of 0. If each session was summarized as a probability distribution over the K = 15 UMAP behaviour clusters, P(k) = fraction of frames assigned to cluster k, renormalized to sum to 1 and given two such probability distributions P and Q, JS divergence is defined as:

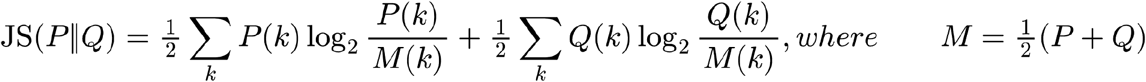

##### Coarse “super-cluster” generation

To identify higher-order classes of behavioral actions, we generated coarse “superclusters” of the UMAP derived finer clusters. To achieve this, we first generated the mean value of all behavior features, on a per cluster basis. Then, we hierarchically clustered the clusters based on their z-scored mean feature values of a subset of 15 non-redundant features using Ward’s method. This yielded 3 superclusters, corresponding to the Male-directed, Female-directed and Non-social behaviors.

#### Distance-based behavior state assignment, and quantification of social preference and choice persistence

For each animal, the body centroid is defined as the arithmetic mean of five midline nodes: nose (primary) and head, neck, trunk and tail-trunk interface (secondary). The Euclidean distance between the subject’s centroid and each conspecific’s centroid was computed per frame and smoothed with a 1 s Gaussian window. Frames in which the smoothed subject-conspecific centroid distance fell below 150 pixels (px) were labelled as close interaction with that conspecific. The 150 px criterion was determined empirically, and the threshold was then held fixed and applied uniformly to every session. Any interval defined as a close interaction that was shorter than 0.5 s were discarded, and consecutive interactions involving the same conspecific separated by less than 2 s were merged into a single epoch. Any interval of at least 2 s containing no interaction epoch with either conspecific was labelled asocial. This procedure divides each session into a sequence of non-overlapping labelled intervals (epochs), each corresponding to a single continuous period of male-directed, female-directed, or asocial behaviour, thus yielding a three-state, mutually exhaustive moment-to-moment description of each session.

##### Social preference

For each session we computed the percentage of total classified time spent in each state, and social preference was expressed as the time in the male-directed state as a fraction of total social time (the summed duration of male- and female-directed epochs), such that 0.5 indicates no preference.

##### Choice persistence

To summarize the temporal structure of state switching, each epoch was represented by one sample per 3 s of its duration (rounded up, so every epoch contributes at least one sample). Samples were concatenated in chronological order and transitions between consecutive samples were counted into a 3 × 3 matrix. The matrix was row-normalized so that each row gives the probability of the next token’s state given the current state. Persistence for each state was taken as the corresponding diagonal element P(i,i); higher values indicate that the animal remained in that state across successive 3 s intervals.

### Neural data analysis

#### Hidden Markov model (HMM) based unsupervised inference of latent neural states

A hidden Markov model (HMM) was used for unsupervised detection of neural state sequences from the population of simultaneously recorded mPFC single units on each day. In the HMM, the population occupies one of *K* hidden states in each time bin, and transitions between states follow a Markov chain with transition matrix *A*. Each state corresponds to a distinct activity pattern across the population of *N* neurons, defined by which neurons tend to spike in it and captured by the emission matrix *E* (*K* × (*N* + 1)), in which each row sums to 1. HMM fitting was adapted from Recanatesi et al. (2022)^47^. Spike trains were segmented into 10 ms bins, and the spiking activity of each neuron was binarized, with 1 indicating that the neuron fired at least once in the bin. Empirically, multiple spikes from one neuron within a time bin were rare (median: 2.2%). In the original implementation, a time bin with multiple neurons firing was reduced to a single neuron drawn at random from the active set. Instead, we split the bin’s contribution equally among all the active neurons. HMMs were fitted with an open-source Python package ssm (https://github.com/lindermanlab/ssm). Model parameters (emission matrix E and transition matrix A) were optimized with the Baum-Welch algorithm, an expectation-maximization (EM) procedure that, given the binarized spike train *Y*, maximizes the log likelihood 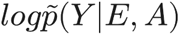 marginalized over hidden state sequences. Iterations stopped when change in log-likelihood was smaller than 10^-10^ or after 500 iterations. Since the procedure only guarantees a local optimum, we ran 60 independent HMM fits on the whole session from random initial conditions, and kept the fit with the highest training log-likelihood for subsequent analyses. The best solution was reproducible: on every session it was reached from multiple independent starts (states agreeing on ≥ 95% of bins). A state was assigned to a bin only if the state’s posterior probability exceeded 0.8 for at least 10 consecutive bins (100 ms). This criterion restricted subsequent analyses to bins with confident state assignment.

##### Choice of the number of HMM states

We chose the number of states K guided by held-out fit. For each session, HMMs with K = 1 - 8 states were fitted with a five-fold cross validation; since neighboring bins are strongly correlated, folds are created with contiguous blocks rather than individual bins, with each block held out exactly once. The held-out log-likelihood per bin, l(K), was computed on the left-out fold. K = 8 was used as the upper bound since the held-out improvement from each added state has become small (from 10.8 x 10^−3^ nats in k =3 to 1.7 x 10^−3^ in k = 7/8). To compare across sessions with different numbers of neurons, we measured gain share (l(k) - l(1) / l(8) - l(1)), the fraction of the improvement over a one-state model at K = 8 that is reached with K states. Most of the gain was reached with a few states: 65% at K= 3, and 80% at K = 4.4. Additional states beyond K = 4 mostly subdivided existing states. Held-out behavior decoding from the HMM states rose from near chance level at K = 1 (0.37, chance 0.33) to 0.63 at K = 4, with smaller gains at higher K (0.72 at K = 8). We therefore used K = 4 which captures most of the held-out gain while keeping the number of latent states closer to the three behavior categories. We also repeated all analyses with K = 3 latent states and observed similar trends.

##### Mapping neural states to behavioral states

Because neural populations differed across recording days, HMMs were fit independently for each session, resulting in session-specific latent states and emission parameters. To enable comparisons across days, we established a mapping between inferred neural states and distance-based behavioral states (Male-directed, Female-directed, and Non-social behaviors). For each session we computed *P*(*n*|*b*), the conditional probability of each neural state given each behavioral state, as the occupancy of that neural state within the frames of that behavioral category, counting only frames with both a defined behavioral label and an admissible neural state (columns normalized to sum to 1). Neural states were then matched to behavioral labels by a global greedy assignment: all (state, behavior) entries of *P*(*n*|*b*) were sorted in descending order and traversed once, accepting each entry only if neither its neural state nor its behavioral state had already been matched. This yields a one-to-one mapping that always assigns the most distinctive state-behavior pairings first, rather than fixing the order of behaviors a priori. This per-session dictionary aligned neural states to a common behavioral reference across days. For K = 4 one neural state necessarily remains unmatched and is retained as a separate, unlabeled state. Additionally, we tested whether the unmatched state represented a finer sub-cluster carved out of a single coarse behavior in which case it can be folded into that behavior or a state spanning several behaviors, which would be misrepresented by any single label. Per session, we computed the distribution of coarse behavioral labels across all frames assigned to the unmatched state, i.e. P(b | n = unmatched), and merged the unmatched state into the behavioral label of the dominant behavior only if that behavior accounted for at least 90% of its frames. Sessions failing this criterion retained the fourth state as its own unlabeled category. The reported effects were all present even when the fourth state was left as its own unlabeled category in all sessions.

##### Neural state accuracy, flicker and entropy

The decoded state sequence was then expanded to a framewise label vector at the video frame rate (60 Hz), with frames falling outside any admissible-state interval (posterior probability < 0.8 sustained for < 100 ms; see above) assigned a null label. Frames carrying no behavioural label were excluded. For each behavioral state b, the neural state accuracy was the proportion of frames within b during which the matched neural state for b was decoded; the session alignment score was the mean of the three per-state values. To quantify rapid transitions in population activity we computed a framewise neural-state flicker metric. Flicker was assigned a value of 1 at time t if the matched neural state changed between t and t+1 while the behavioral state remained unchanged, and 0 otherwise (the final frame was set to 0 by construction); neural transitions coincident with a behavioral transition are expected and were therefore not counted as flicker. Flicker rate was expressed per behavior as the percentage of frames within that behavior scored as flicker. To focus on how persistently a neural state is maintained within a behavior, rather than in how states are updated at behavioral boundaries, all flicker analyses were repeated with a more stringent temporal exclusion, in which every frame falling within ±1 s (±60 frames) of a coarse behavior change point (male-directed / female-directed / non-social) was treated as ineligible and dropped from both the numerator and the denominator of the flicker rate, and we observed similar effects. To quantify how distributed population activity was across latent states within a behavior, we computed the Shannon entropy of the neural-state distribution conditioned on each behavioral state, *H*(*b*) =− Σ *P*(*n*|*b*) *log*2 *P*(*n*|*b*), in bits. Session entropy was reported as the mean of the three per-behavior values.

#### Linear discriminant analysis (LDA) and Support Vector Machine (SVM) based discriminability and decoding of behavior states

For Linear discriminant analysis (LDA), spike counts from each unit were binned in 0.5 s bins across the behavioral session. For each time bin, binned counts from all remaining units were then assembled into a population activity vector and assigned a behavioral label (male-directed, female-directed, or asocial) based on the concurrent behavioral state. Units firing fewer than 100 spikes across the labeled bins were excluded. Such units contribute negligible within-class variance, which both renders the within-class scatter matrix ill-conditioned and, because LDA maximizes a between- to within-class variance ratio, allows chance spike placement in near-silent units to dominate the discriminant axes. LDA was applied to all labeled bins, yielding two discriminants (number of classes − 1), and for each behavioral class, the center and dispersion in discriminant space were estimated using the minimum covariance determinant estimator (robustcov, outlier fraction 0.1) and class distributions were summarized with 95% probability ellipses. Separability was quantified as the pairwise Euclidean distance between robust class centers in the two-dimensional discriminant space; Mahalanobis distances, normalized by the pooled robust covariance of each class pair, yielded similar results. To control for class imbalance, analyses were repeated after subsampling male- and female-directed bins to the smaller of the two counts (and capping non-social bins at that count), with qualitatively similar results.

Decoding was assessed with a nested LDA-SVM classifier under 5-fold cross-validation. Folds were stratified by behavioral state, preserving class proportions across folds. Within each fold, LDA axes were estimated from the training bins only, and both training and held-out bins were projected onto those axes; a multiclass linear Support Vector Machine (SVM) was then fitted to the projected training bins and used to predict the labels of the held-out bins. Decoding accuracy was the fraction of correctly classified held-out bins, averaged across folds.

#### Effective dimensionality and manifold geometry

Spike counts were binned in 0.5 s bins across the behavioral session for each unit, yielding a units × time-bin matrix in which each bin is a point in the N-dimensional neural state space defined by the activity of the N simultaneously recorded mPFC neurons. Each bin was labeled male-directed, female-directed or asocial according to the concurrent behavioral state; bins not assignable to a state were discarded. The set of bins sharing a label constitutes that state’s manifold object. For each manifold object separately, principal component analysis was applied to the mean-centered sub-matrix of bins belonging to that state, and effective dimensionality was quantified using the participation ratio of the resulting eigenvalue spectrum:

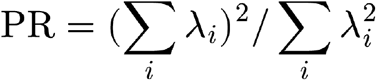

where *λ*_i_ are the eigenvalues of the within-state covariance matrix. PR is a continuous measure of how evenly variance is distributed across dimensions, equalling 1 when a single direction captures all variance within the object and N when all directions contribute equally; Effective dimensionality is reported as PR normalized by the number of units entering the estimate (PR/N), which expresses it as a fraction of the maximum dimensionality attainable for that population size and bounds the measure in (0, 1].

This normalization does not correct for recording yield: PR grows sub-linearly with the number of units, so PR/N retains a dependence on N. Comparability across recording days was therefore enforced by holding the number of units constant across sessions within each mouse. On each of 250 iterations, n units were drawn without replacement from that day’s population, where n = 85% of the minimum yield across that mouse’s days. Sampling was without replacement because duplicated units are perfectly correlated and inflate the leading eigenvalues, biasing PR downward; the 0.85 factor ensures that even the lowest-yield day draws a genuinely different subset on each iteration rather than returning its full population every time. All three manifold objects were recomputed from the same unit subsample on each iteration, so within-session comparisons across states are matched by construction. Days with fewer than 50 units, or fewer than 60% of the median yield for that mouse, were flagged and considered for exclusion, since one low-yield day otherwise caps n for every other day in that mouse.

#### Single-unit tuning and lifetime sparseness

For each unit and each behavioral state, spike rasters were extracted in a short ±3 s window around the onset of each behavioral state, since transition between states happen in a spontaneous and sometimes fast timescales. The peri-state time histogram (PSTH) was computed by averaging across trials at 1 ms resolution and converting to spike rate (spikes/s). The PSTH was then smoothed using a Gaussian kernel (500 ms window) to obtain a smoothed firing rate estimate r(t). The baseline firing rate for each unit was estimated from the first half of the pre-onset window, spanning −3 to −1.5 s relative to state onset, and for each state j, the smoothed PSTH was then normalized to the baseline mean firing rate to yield a fold-change timeseries:

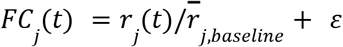

where *r̅_j_*_,*baseline*_ is the mean firing rate over the baseline period for state j and *ε* is a small constant (machine epsilon) to prevent division by zero for silent neurons. The response magnitude to each state was computed as the mean fold-change over a −0.5 to +3 s window centered on state onset. This window captures both the immediate onset response and sustained activity in the first 3 s of the state, while the 0.5 s pre-onset inclusion accommodates short anticipatory ramps. Modulation to behavioral states was assessed using the peak absolute z-score of the smoothed PSTH within the −0.5 to +3 s window around state onset relative to the baseline period).

##### Triangle plots

The three state response magnitudes [*r̂_M_*, *r̂_F_*, *r̂_NS_*] were L1-normalized across states:

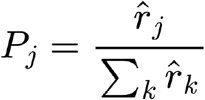

such that 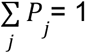 and each *P_j_* represents the fraction of the total response allocated to state j. For visualization of this response selectivity, each unit is represented on an equilateral triangle with vertices at (0,0,1), (0,1,0), and (1,0,0), where the three axes correspond to the normalized responses to Male, Female, and Non-social behaviors. Units are projected onto the plane formed by the triangle’s intersections with the Cartesian axes^88^. On each day a unit was assigned a behavioral-state tuning label by winner-take-all selection over the baseline-normalized, L1-normalized response to the three social states (M / F / NS), gated on a peak |z| ≥ 3 response relative to the pre-transition baseline; units failing the responsiveness gate were labelled untuned, and excluded from Lifetime Sparseness computation described below.

##### Lifetime sparseness(LS)

LS was computed from the normalized response vector P = [P_M_, P_F_,P_NS_] using the bias-corrected formula:

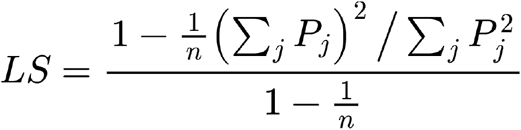

where n = 3 (number of behavioral states). LS ranges from 0 (equal response to all states; broadly tuned) to 1 (response to a single state; maximally selectively tuned). The bias-correction factor (1 − 1/n) in the denominator rescales the index to span the full [0, 1] range for any n. Modulated units were assigned to be tuned to a preferred state corresponding to the state with the highest normalized fold-change response (argmax P_j_).

#### Single-unit intrinsic timescales

We estimated intrinsic timescales of single-neuron activity from the spike-count autocorrelation computed during the 10-min resting-state period at the beginning of each session. Spontaneous firing rates were computed from this same window. Spike trains were binned into 5 ms bins across this window, yielding a spike-count sequence A(t_1_…t_N_) per unit with N bins. For each unit the normalised autocorrelation was computed at lags t_j_ = 0 to 5 s as

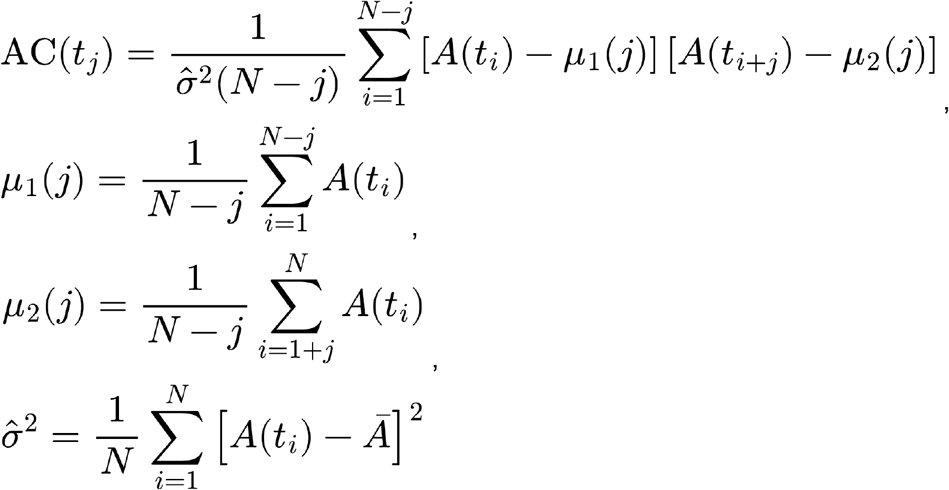

where μ₁(j) = mean of the leading segment A(1:N−j), μ₂(j) = mean of the lagging segment A(1+j:N), and 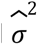 is the population variance of the full binned trace.

##### Autocorrelation fits

The autocorrelation was truncated at 5 s to avoid fitting long-lag noise. The refractory period and shot noise produce a dip immediately after lag 0 followed by a rebound, hence the fit was started at the end of the rebound, identified in two stages: The refractory dip was taken as the first local minimum of AC within 50 ms of lag 0, and the start of the decay was taken as the maximum of the autocorrelation over the 500 ms following the dip, after smoothing with a 5-bin moving average to the post-dip segment.

We fitted the autocorrelation with linear mixtures of exponential decay functions as described previously^60^:

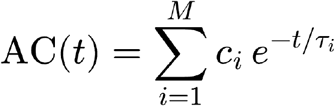

Here *τ*_i_ are the timescales, *c*_i_ are their mixing coefficients, and M is the number of distinct timescales. We fitted models with M ranging from 1 (single timescale) up to 4 (four timescales) and selected the best-fitting model using the Bayesian information criterion (BIC), among the candidate orders in which every component contributed ≥1% of the total amplitude ∑ *c*_i_. Only the units for which the model provided a good fit (coefficient of determination R^2^ > 0.5) were included in subsequent timescale analyses. For these neurons, the effective timescale was defined as the average of individual timescales *τ*_i_ weighted by their mixing coefficients *c*:

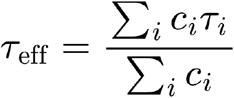

The effective timescale captures the overall temporal persistence of neural activity, integrating over the multiple timescales that contribute to a neuron’s dynamics. Since session-level timescales are summarized as a median across units, sessions with low yield in which fewer than ten units satisfied the fit-quality criteria were excluded from session-level comparisons. Fitting the activity of each neuron using a single-exponential fit without applying any selection criteria to the goodness of fit also yielded qualitatively similar results.

#### Tuning drift and comparisons across hormonal states for tracked units

A unit entered the tuning drift analysis only if it was tracked on ≥ 2 low-E2 days and ≥ 2 high-E2 days, so that within-state stability could be estimated in both hormonal states from the same unit. Within each hormonal state, tuning drift was quantified over chronologically consecutive tracked-day pairs as the proportion of pairs on which the unit’s tuning label changed. Across states, each unit was summarized by a single representative label per state (modal label within low E2 and high E2 state; ties broken by the last day in that state). We stratified tracked units by their baseline selectivity, taken as the mean LS across that unit’s low-E2 days, using a threshold of 0.25. The stratification was defined on low-E2 days alone and the same units were then followed into high E2. Units with high baseline LS have a well-separated preferred state, so their label is expected to be reproducible across days in either hormonal state and they offer little dynamic range in which a change in stability could be detected; The analysis therefore focused on the low-LS bucket, units that respond more broadly across the three states under low E2, where we asked whether these broadly responsive units instead acquire a more reproducible and stable state preference under high E2.

For all other metrics such as LS, intrinsic timescale, spontaneous activity, a unit was included if it was tracked on at least one low-E2 day and at least one high-E2 day; Each unit’s values were then pooled within each state by taking a central-tendency statistic across its low-E2 days and across its high-E2 days, and the two per-unit values were compared.

#### Functional connectivity during resting-state

Cross-covariance was computed for all unique pairs of simultaneously recorded units within each session. For each pair, the raw cross-correlation was computed using MATLAB’s xcorr function: *C*(*τ*) = *xcorr*(*a*, *b*, *lag*) where a and b are the column-vector spike trains of the two units, and lag = 25 ms defines the maximum temporal offset evaluated. The raw cross-correlation was normalized to a cross-covariance estimate by removing the expected correlation as described previously^89^:

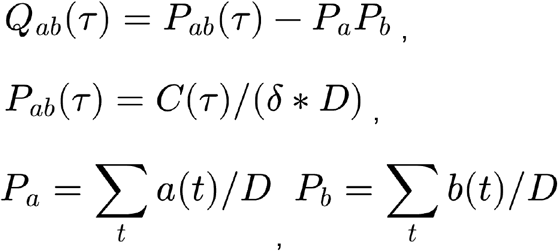

where *P_ab_*(*τ*) is the normalized cross-correlation, *P_a_ and P_b_* are the mean firing rates of the two units (spikes per bin), *δ* = 1 ms is the bin width and D = total number of time bins. Subtracting *P_a_P_b_* removes the baseline coincidence rate expected from two independent Poisson processes with the same mean firing rates, leaving *Q_ab_*(*τ*) as a measure of excess synchrony or suppression at each lag. The raw *Q_ab_* was then smoothed with a Gaussian kernel of width 4 ms to reduce noise before significance testing. A theoretical confidence interval for *Q_ab_* under the null hypothesis of independence was derived analytically based on the standard deviation of the cross-covariance estimate for two independent Poisson spike trains:

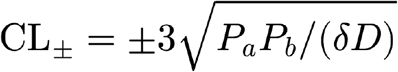

Each pair was classified as significantly positively or negatively correlated if the cross-covariance exceeds CL+ or CL-respectively within a ±5 ms window. Pairs whose cross-covariance peak occurred at zero lag were excluded from further analyses, since synchronous peaks are more consistent with shared common input than with a directed interaction between two units. Further, sessions in which single-unit yield was insufficient (< 60) to estimate a network with these constraints were excluded.

##### Network graph visualization

For each recording day, the connectivity matrix separately for positive and negative cross-covariance was passed to MATLAB’s graph function using the ‘upper’ flag to construct an undirected graph. A standard force-directed layout was used placing highly connected hubs centrally.

##### Normalized node degree

For each recording day, the degree of each node was normalized and calculated as Normalized degree = Number of direct significant connections / (N − 1) where N is the total number of nodes in the network. This normalization scales degree to [0, 1], where 1 represents a unit connected to every other unit.

##### Chord-style plots

For each recording day, an individual-unit chord diagram was generated from the same connectivity matrix. Each such unit was represented as a node on a unit circle, equally spaced within its tuning group and ordered by group (male-, female-, and self-tuned) with a fixed angular gap separating groups. Only tuned units surviving the lifetime-sparseness criterion (LS > 0.05) were included. Every significant pair contributed a chord, drawn as a quadratic Bézier curve bowed toward the center.

##### Connection probability as a function of signal correlation

For every simultaneously recorded pair that entered the cross-covariance analysis (positive and negative connections were analysed separately), a signal correlation was computed as the Pearson correlation coefficient between the two units’ session-long firing-rate time courses, binned at 1 s over the whole behavioural session. For visualization, connection probability was modelled as a function of signal correlation using logistic regression with signal correlation as the sole predictor. As a model-free complement, the pooled distribution of signal correlations was split at its 25th and 75th percentiles, and the mean connection probability was computed within the lowest (Q1) and highest (Q4) quartile separately for each recording day and for positive and negative connections. Quartile thresholds were defined once on the pooled data so that the same signal-correlation bounds applied to every day.

### Mathematical modeling

#### Energy landscape model and attractor well depths

To develop a simple conceptual model for how estrogen reconfigures the dynamical landscape governing neural-state transitions, we constructed a phenomenological energy landscape model constrained by empirical state dwell times. Under Kramers’ escape-rate theory^52^, the mean dwell time in a metastable state relates to the depth of its basin as *τ_dwell_* = *τ*_0_. *e*_(Δ*U*/*D*)_, where ΔU is the depth of the basin, D scales the stochastic fluctuations, and *τ* is a constant with units of time set by the local dynamics within the basin. Rearranging, well depth is estimated as Δ*U* = *D* · [*log*(*τ_dwell_*) − *log*(*τ*_0_)]. We set D = 1 and *τ*_0_ = 1, and a single well depth has no interpretation on its own; only low-versus-high-E2 differences in ΔU are meaningful.

Dwell times were taken from the HMM neural states, restricted to episodes of a neural state occurring while the matching behavior was expressed and beginning within 10 s of a matching behavior epoch onset but episodes extending beyond that window retained their full duration. This restriction was applied because unrestricted matched dwell times could partly re-express any difference in behavior epoch duration as a difference in neural persistence and anchoring to a fixed window after epoch onset samples each epoch at a comparable phase of the interaction. Well depths were computed separately for male-directed (M), female-directed (F), and non-social (NS) states, from the median dwell time within each session (since the dwell distributions are strongly right-skewed) and then averaged.

For visualisation, we constructed a three-well potential over a two-dimensional state space, s = (s₁, s₂) ∈ [0,1]², with the male-directed, female-directed, and asocial attractors placed at [0.8, 0.2], [0.2, 0.8], and [0.2, 0.2] respectively. Each attractor was represented as an isotropic Gaussian well whose depth is the empirically derived ΔU for that state and whose width σ = 0.12 was chosen to be narrow enough that neighbouring wells do not overlap appreciably at the saddles between them:

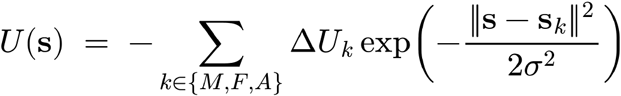

This landscape is primarily a visualisation tool within a dynamical systems framework where metastable states are modeled as attractors of an effective dynamical system, with transitions between states occurring when fluctuations drive population activity out of one basin of attraction and into another^49,50,52,90^.

#### Mean-field dynamical systems model

To test whether the measured change in recurrent coupling, imposed on a circuit was sufficient to produce the observed changes in neural dynamics, we implemented a dynamical systems model. The circuit architecture was chosen to be the minimal circuit consistent with the structure of the behaviour and of the measured connectivity. The smallest circuit motif that generates mutually exclusive, self-sustaining states is a pair of populations, each recurrently exciting itself and inhibiting the other, which yields winner-take-all bistability. This motif also matches the organisation recovered from the cross-covariance graphs. The two model populations were therefore identified with male and female tuned ensembles, and the recurrent weight was set from the within-cluster connection probability in each hormonal state. We use a “mean-field” or a “rate” approximation where we qualitatively capture the behavior of the network by describing it at each time point, in terms of the average activity of all the male and female-tuned neurons, which are represented by the two nodes. Population activity, *x* (the male-preferring population) and *x* (the female-preferring population), evolved according to the following firing-rate equations^12,13^:

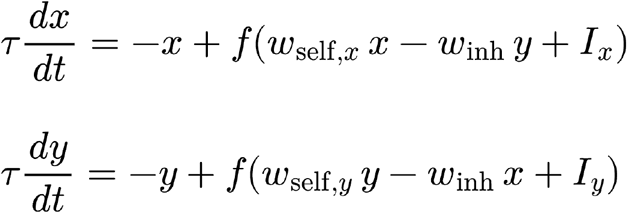

where *x*, *y* ∈ [0 1] are normalized population firing rates, *w_self_* is the within-population recurrent excitation weight, *w_in_*_ℎ_ is the cross-population inhibition weight, and the excitatory and inhibitory inputs are approximated to combine linearly. *I_x_*, *I_y_* are constant external inputs to the two populations (we simulate the dynamics at a short timescale where we assume constant external drive). The population time constant *τ* (at which activity would decay without any input) is set to 100 ms. The input-output transfer function f that models the non-linear activation function is a bounded, saturating sigmoid:

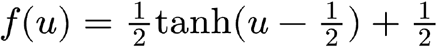

which maps an unbounded net synaptic input *u* onto the open interval (0, 1), and yields f(0) ≈ 0.27 i.e. a population that is largely but not completely quiescent at rest. To model male-interaction epochs, a small asymmetric external drive was applied to bias the system toward the male-preferring population:

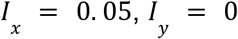

##### Parameter grounding in the functional-connectivity data

The recurrent weight *w_self_* was set from the measured mean within-ensemble connection probability (CP) of the positive cross-covariance graphs during low and high E2. Because CP is a dimensionless probability and *w_self_* is a synaptic weight, the two were related by a single fixed scaling factor *k*:

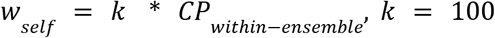

*k* was chosen such that low-E2 operating point sits just inside the bistable regime of the model and is held identical across hormone states. Since the male-preferring and female-preferring within-cluster CPs were closely matched, the model was run in symmetric configuration, with both populations assigned the same *w_self_*

Since the between-cluster connection probability of the negative cross-covariance graph is an order of magnitude below the positive within-cluster CP, and negative and positive cross-covariances are not detected with comparable sensitivity, and their magnitudes cannot be placed on a common scale. *w_in_*_ℎ_ was therefore treated as a free parameter and fixed at *w_in_*_ℎ_ = 1. 8 for both hormone states, a value that places the low-E2 operating point inside the bistable regime. Holding *w_in_*_ℎ_ fixed across states was also consistent with the negative between-cluster CP showing no significant change during high E2.

##### Phase-plane analysis

For each hormone state, the vector field 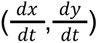 was evaluated on a 50 × 50 grid, and the x- and y-nullclines extracted as the zero contours of 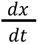 and 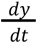 respectively. Flow direction is shown as a quiver field, and flow magnitude 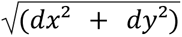 is overlaid as a filled contour map or shown by the arrow size.

##### Stochastic simulations

State dynamics were simulated by adding noise to the deterministic flow. Trajectories were integrated by forward Euler with dt = 0.5 ms for T = 500 ms. Independent Gaussian perturbations of standard deviation *σ* = 0.05 were added to x and y every fifth integration step (every 2.5 ms) and the state was clipped to [0, 1] after every update. At every time point the system was assigned to one of three states using a fixed margin around the x = y diagonal (margin = 0.05): x-dominant (male neural state) if x > y + 0.05; y-dominant (female neural state) if y > x + 0.05; transition state otherwise; Each trial was initialized in the x-dominant basin at x(0) = 0.8 + 0.05*ξ, y(0) = 0.2 + 0.05*ξ with ξ ∼ N(0, 1), so that every trial begins in the male-preferring state and any transition to the female-preferring state is a noise-driven escape. Ten trials per hormone state were simulated for phase-plane trajectory display and 100 independent trials per hormone state for the dwell-time statistics.

##### One-dimensional energy landscape

To compare the model’s prediction for how estradiol reshapes the landscape against the empirically estimated energy landscape (derived from HMM dwell times), the two-dimensional flow was projected onto a single decision axis. This yields a one-dimensional pseudo-potential whose well depth sets the barrier separating the two behavioural states, similar to the quantity estimated empirically from the HMM dwell-time distributions. The system is evaluated along the anti-diagonal y = 1 − x, the axis along with the two populations trade activity. At each point on this line we took the component of the deterministic flow pointing along it, and integrated it outward from the saddle to obtain a potential, U. Flow directed away from the saddle therefore lowers U, placing the saddle at zero and each attractor at the bottom of a negative well. Well depth ΔU is the barrier the system must cross to switch states, and is the model’s counterpart to the empirical landscape.

#### Inferential statistical analysis

For all statistical tests, p values were compared against an *α* significance threshold of 0.05. Where necessary, we used a linear mixed-effects model (LME) to account for the repeated measures within animals by modeling animal identity as a random effect. Correction for multiple comparisons was applied within each reported family of tests (for a set of measurements presented together in a single panel), using the Benjamini-Hochberg false discovery rate (FDR) method. LMEs were built using the MATLAB function ‘fitlme’ with ‘FitMethod’ set to ‘Maximum Likelihood’.

## Notes

### Competing Interest Statement

The authors have declared no competing interest.

